# Hierarchical Complexity of the Adult Human Structural Connectome

**DOI:** 10.1101/389569

**Authors:** Keith Smith, Mark E. Bastin, Simon R. Cox, Maria C. Valdés Hernández, Stewart Wiseman, Javier Escudero, Catherine Sudlow

**Affiliations:** Usher Institute for Population Health Science and Informatics, Medical School, University of Edinburgh, Edinburgh, EH16 4UX, UK; Centre for Clinical Brain Sciences, Western General Hospital, University of Edinburgh, Edinburgh, EH4 2XU, UK; Centre for Cognitive Ageing and Cognitive Epidemiology, Department of Psychology, University of Edinburgh, Edinburgh, EH8 9JZ, UK; Row Fogo Centre into Ageing and the Brain, Edinburgh Dementia Research Institute, University of Edinburgh, Edinburgh, EH16 4SB, UK; School of Engineering, Institute for Digital Communications, University of Edinburgh, Edinburgh, EH9 3FB, UK

## Abstract

The structural network of the human brain has a rich topology which many have sought to characterise using standard network science measures and concepts. However, this characterisation remains incomplete and the non-obvious features of this topology have largely confounded attempts towards comprehensive constructive modelling. This calls for new perspectives. Hierarchical complexity is an emerging paradigm of complex network topology based on the observation that complex systems are composed of hierarchies within which the roles of hierarchically equivalent nodes display highly variable connectivity patterns. Here we test the hierarchical complexity of the human structural connectomes of a group of seventy-nine healthy adults. Binary connectomes are found to be more hierarchically complex than three benchmark random network models. This provides a new key description of brain structure, revealing a rich diversity of connectivity patterns within hierarchically equivalent nodes. Dividing the connectomes into four tiers based on degree magnitudes indicates that the most complex nodes are neither those with the highest nor lowest degrees but are instead found in the middle tiers. Spatial mapping of the brain regions in each hierarchical tier reveals consistency with the current anatomical, functional and neuropsychological knowledge of the human brain. The most complex tier (Tier 3) involves regions believed to bridge high-order cognitive (Tier 1) and low-order sensorimotor processing (Tier 2). We then show that such diversity of connectivity patterns aligns with the diversity of functional roles played out across the brain, demonstrating that hierarchical complexity can characterise functional diversity strictly from the network topology.

## Introduction

The physical connections between regions of the human brain transcend their geometrical localities to support globally efficient and complex functional principles^1,2^. Characterisations of this structure as a network allows us to probe hidden architectural patterns, facilitating a deeper understanding of the brain’s wholescale organisation^3–5^. Such characterisations are enabled by network indices, which are used to measure and rank specific topological properties of networks, and null models, which are used to compare how networks differ from random networks designed with in-built topological characteristics. Yet much remains to be understood about these patterns and how they support the brain’s multifaceted roles in, for example, information processing, creativity and cognition. Global network characteristics are modelled on the basis of brain regions and the connections between them. Important findings show efficiency^3^, fractal modular organisation^4^ and rich-club structures between hub nodes^6^ of brain networks. It has also been suggested that the inability to find simple generative models of the connectome implies the existence of a variety of different biological mechanisms working in conjunction with each other^7^. Methods to combine such mechanisms to explain brain structure have been attempted with moderate success, suggesting distance-based penalties and a tendency for neighbouring nodes to share neighbours being two key factors^8,9^. However, it has yet to be determined whether dissimilarity of connectivity patterns is itself a feature which can advance our understanding of brain structure. Just such a feature can be extracted using the recently developed hierarchical complexity paradigm^10,11^.

Here, for the first time, we analyse the hierarchical complexity of the human structural connectome created from structural and diffusion magnetic resonance imaging (MRI) data acquired from a sample of working age individuals. This includes a detailed analysis of complexity within the hierarchical levels of these connectomes. This recently introduced paradigm has been validated only in an electroencephalogram (EEG) functional connectivity study^10^. It posits that network complexity is characterised by nodes of the same degree (hierarchically equivalent nodes) being connected in highly variable ways with respect to the degrees of nodes they connect to (having highly variable connectivity patterns), as illustrated in Figure 1. This concerns wholly separate considerations of topology to the well-known paradigms of small-world^12^ and scale-free^13^ complex networks— the former stemming from the idea that there are no more than six degrees of separation between any two people, which is exhibited with network characteristics of high clustering and low shortest path lengths, and the latter being that complex networks display power-law degree distributions, crudely identifiable by having few hub nodes with many connections and many peripheral nodes with few connections. Similarly, in seeking to define the notion of complexity, it takes a different stance to the standard notion of complexity arising between random and ordered systems^14^, instead proposing that both such systems have inherently more predictable connectivity patterns than real-world complex networks^10^.

**Figure 1.**
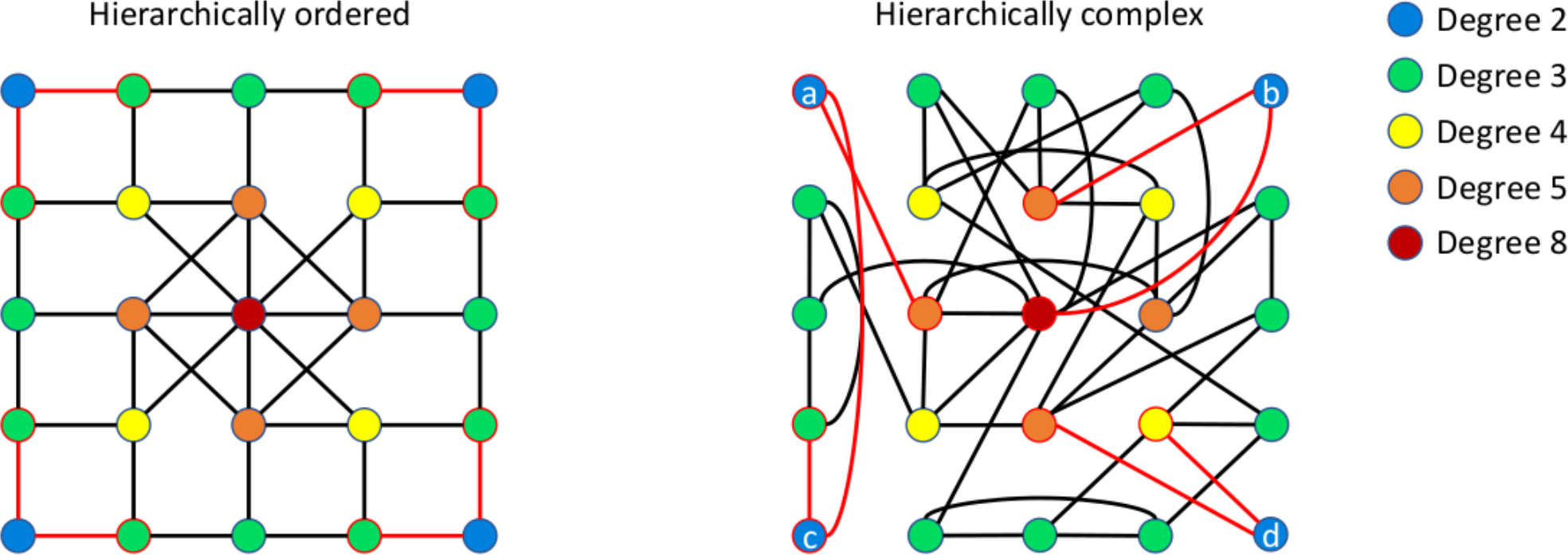
Illustration of hierarchical complexity. Two networks are shown with 25 nodes, 44 edges and identical degree distributions. Node colours signify degrees as in the legend. The connectivity patterns (degrees of nodes a node is connected to) of degree 2 nodes are highlighted in the images by red edges and node boundaries. In a hierarchically ordered network, left, same degree nodes have homogeneous connectivity patterns. In a hierarchically complex network, right, same degree nodes have heterogeneous connectivity patterns. For example, node c is connected to only low degree nodes (2 and 3), and node b to only high degree nodes (5 and 8).

In the human brain, we particularly expect such behaviour. The brain is composed of numerous regions with myriad functional specialisations, a phenomenon which we hypothesise to necessitate a wide variety of connectivity patterns in the supporting structure. We test this hypothesis by comparing the network index for hierarchical complexity of structural connectomes against those of three node- and edge-matched randomised models. Complementing this, we seek to answer where in the network hierarchy, as well as in the brain biology, such complexity is prominent. We do this by splitting the structural connectome into hierarchical tiers and performing within tier analyses before analysing which regions lie consistently (in more than two thirds of participants) within one of these tiers. Critically, it is well established that hub regions exist in the brain^15,16^ and it is suggested that their degradation is a key mechanism in brain disorders^17^. Therefore, it is of interest to understand whether or not hub nodes are drivers behind the brain’s structural complexity or if other hierarchy levels are more complex and what implications this may have in our understanding of brain connectivity and how this could be implemented to aid our understanding of pathology. Finally, we study specific notable ROIs to understand how their unique connectivity patterns may be explained by their known functional roles.

## Materials and Methods

For reference, a block diagram of the methodological pipeline used in this study is provided in Figure 2. Two sets of network analyses were produced to undertake a comprehensive analysis of hierarchical complexity in the adult human structural connectome. The first concerned the hierarchical complexity of binary structural connectomes. The second concerned the hierarchical tiers most responsible for the hierarchical complexity of the binary connectomes and the regions within these tiers. The latter was then used to compute ROI connectivity profiles, relaying the fractions of Tier 1, Tier 2, Tier 3 and Tier 4 nodes to which the ROI was connected.

**Figure 2.**
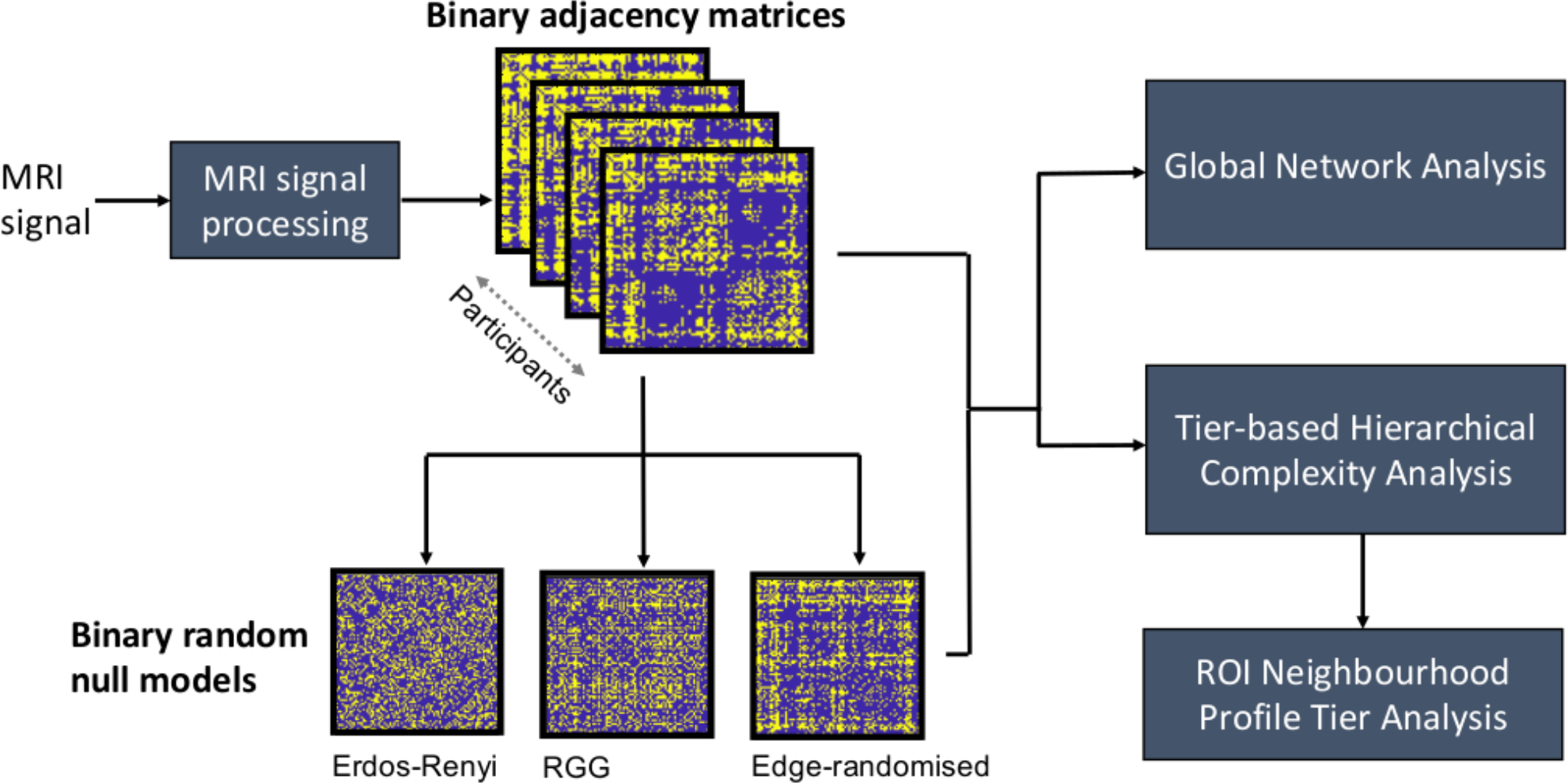
Block diagram of the employed methodological pipeline. Adjacency matrices are computed from the MRI signal. Random models are then generated with matching network size and density. Network indices are then computed from which the results are derived.

### Subjects

Eighty normal, healthy volunteers (40 males, 40 females) aged 25–64 (median 43, IQR 17) years were recruited by advertisement from staff working at the University of Edinburgh, the Western General Hospital and Royal Infirmary, Edinburgh, Scotland. Health status was assessed using medical questionnaires and all structural MRI scans were reported by a neuroradiologist.

Volunteers were recruited if they were native English speakers, were not on any long-term medication, had not been diagnosed with any chronic medical condition including diabetes mellitus or hypertension, had not undergone previous cranial surgery, and were able to undergo brain MRI. The study was approved by the Lothian Research Ethics Committee (05/S1104/45), and subjects gave written informed consent.

### MRI acquisition

All MRI data were acquired using a GE Signa Horizon HDxt 1.5 T scanner (General Electric, Milwaukee, WI, USA) using a self-shielding gradient set with maximum gradient strength of 33 *mT*/*m* and an 8-channel phased-array head coil. Briefly, subjects provided high resolution structural (*T*_1_ -, *T*_2_ -, *T*_2_* - and fluid attenuated inversion recovery (FLAIR)-weighted scans) and diffusion MRI data in the same session. The diffusion MRI examination consisted of 7 *T*_2_- weighted (*b* = 0 *s mm*^−2^) and sets of diffusion-weighted (*b* = 1000 *s mm*^−2^) single-shot spin-echo echo-planar (EP) volumes acquired with diffusion gradients applied in 64 non-collinear directions^18^. Volumes were acquired in the axial plane with a field-of-view of 256 × 256 *mm*, contiguous slice locations, and image matrix and slice thickness designed to give 2 *mm* isotropic voxels. A 3D *T*_1_-weighted inversion recovery-prepared fast spoiled gradient-echo (FSPGR) volume was also acquired in the coronal plane with 160 contiguous slices and 1.3 *mm*^3^ voxel dimensions.

### Image processing

Each 3D *T*_1_-weighted FSPGR volume was parcellated into 85 cortical (34 per hemisphere) and sub-cortical (eight per hemisphere) regions-of-interest (ROI), plus the brain stem, using the Desikan-Killiany atlas and default settings in FreeSurfer v5.3 (http://surfer.nmr.mgh.harvard.edu). The results of the segmentation procedure were visually checked for gross errors and then used to construct grey and white matter masks for use in network construction and to constrain the tractography output. Using tools provided by the FDT package in FSL (http://fsl.fmrib.ox.ac.uk/fsl), the diffusion MRI data were pre-processed to reduce systematic imaging distortions and bulk subject motion artifacts by affine registration of all subsequent EP volumes to the first *T*_2_-weighted EP volume^19^. Brain extraction was performed on the registered *T*_2_-weighted EP volumes and applied to the fractional anisotropy (FA) volume calculated by DTIFIT in each subject^20^. The neuroanatomical ROIs determined by FreeSurfer were then aligned from 3D *T*_1_-weighted volume to diffusion space using a cross-modal nonlinear registration method. As a first step, linear registration was used to initialize the alignment of each brain-extracted FA volume to the corresponding FreeSurfer extracted 3D *T*_1_-weighted brain volume using a mutual information cost function and an affine transform with 12 degrees of freedom^19^. Following this initialization, a nonlinear deformation field based method (FNIRT) was used to refine local alignment^21^. FreeSurfer segmentations and anatomical labels were then aligned to diffusion space using nearest neighbour interpolation.

### Tractography

Whole-brain probabilistic tractography was performed using FSL’s BedpostX/ProbTrackX algorithm^22^. Probability density functions, which describe the uncertainty in the principal directions of diffusion, were computed with a two-fibre model per voxel^23^. Streamlines were then constructed by sampling from these distributions during tracking using 100 Markov Chain Monte Carlo iterations with a fixed step size of 0.5 *mm* between successive points. Tracking was initiated from all white matter voxels and streamlines were constructed in two collinear directions until terminated by the following stopping criteria designed to minimize the amount of anatomically implausible streamlines: (i) exceeding a curvature threshold of 70 degrees; (ii) entering a voxel with FA below 0.1; (iii) entering an extra-cerebral voxel; (iv) exceeding 200 *mm* in length; and (v) exceeding a distance ratio metric of 10. The distance ratio metric^24^ excludes implausibly tortuous streamlines. For instance, a streamline with a total path length 10 times longer than the distance between end points was considered to be invalid. The values of the curvature, anisotropy and distance ratio metric constraints were set empirically and informed by visual assessment of the resulting streamlines. Data is available online at https://www.brainsimagebank.ac.uk.

### Network construction

Connections were recorded in an 85 × 85 adjacency matrix, where the entry *a*_*ij*_ denotes the connection (edge) weight between node *i* and node *j*, where each node represents the aggregated tissue of one of the 85 ROIs. FA-weighted networks were computed by recording the mean FA value along interconnecting streamlines. Across the cohort, only connections which occurred in at least two-thirds of subjects were retained^25^. Self-connections were removed as these cause unwanted complications to network analyses, and if no streamlines were found between a pair of nodes, the corresponding matrix entry was set to zero. Network construction failed in one subject giving structural connectome data for seventy-nine subjects.

### Hierarchical complexity

The hierarchical complexity of the structural connectomes was implemented to analyse how similar the connections established by nodes of the same degree were in terms of the degrees of the nodes they were connected to. This was achieved by computing the variability of the ordered node neighbourhood degree sequences. Let *G* = (𝒱, ε) be a graph with node set 𝒱 = {1, …, *n*} and edge set ε = {(*i*, *j*): *i*, *j* ∈ 𝒱}, and let 𝒦 = {*k*_1_, ⋯, *k*_*n*_} be the set of degrees of *G*, where *k*_*i*_ is the number edges adjacent to vertex *i*. Further, let 𝒦_*p*_ be the set of nodes of degree *p*. For neighbourhood degree sequence **s**^*p*^_*i*_ = {*s*^*p*^_*i*_(1), ⋯, *s*^*p*^_*i*_ (*p*)} of node *i* of degree *p*, the hierarchical complexity is

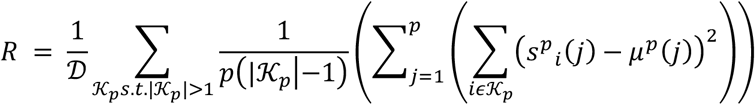

where 𝒟 is the number of distinct degrees in the network and *μ*^*p*^(*j*) is the mean of the *j*th entries of all *p* length neighbourhood degree sequences^10^. For the tier-based analyses, we used degree specific hierarchical complexity by averaging hierarchical complexity over a given range of degrees, i.e. the degrees within the given tier definitions.

### Connectivity and network analysis

Connectivity matrices were first binarised by setting all non-zero entries to 1 to obtain binary network topologies. For each connectivity matrix, three randomised graphs were generated with the same number of nodes (always *n* = 85) and edges (*m* = 1281.5 ± 136.72).

Firstly, Erdos-Renyi random graphs were generated as a baseline randomisation in which each node has an equal probability of being connected to any other. Random uniform weights were computed for each edge and the *m* largest weights were kept as edges. Secondly, to test the differences between brain connectivity and a closest distance-based connectivity of points placed randomly in 3D space, we generated random geometric graphs (RGGs)^26^. These were created by generating uniformly random 3D coordinates (representing nodes) and computing the distances between each pair of coordinates (representing weighted edges). The *m* closest number of edges were then taken as the binary topology. Thirdly, we tested the difference between brain connectivity and graphs with the same degree distribution but with randomised edges, known as configuration models^27^. This allowed us to test the hierarchical complexity of the human structural connectome against a randomised null model controlling for graph heterogeneity and thus overcame any bias found simply due to a different level of heterogeneity in graph degrees.

Hierarchical complexity was computed for all connectomes and null models alongside the following network indices:

1. The degree variance^28^,

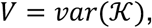

characterising the spread of the degree hierarchy and thus associated with the dominance of the network hubs.
2. Assortativity^29^,

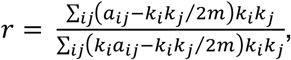

characterising the similarity of neighbouring node degrees and thus helping to understand whether or not hierarchically equivalent nodes group together.
3. The normalised clustering coefficient^10,12^,

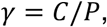

where *P* = 2*m*/*n*(*n* − 1) is the network density (number of edges out of total possible in a network with *n* nodes) and 𝐶 is the global clustering coefficient defined as the ratio of the number of triangles (3 nodes all sharing edges) in the network and the number of triples (paths of length 2) in the network. This *measures the extent of segregation within the network, i.e. the tendency for nodes to cluster into highly intra-connected groups*. These other indices are computed for comparisons and to allow for greater insight into topological differences.

### Hierarchy tiers

Once the global connectivity patterns were assessed we then performed a more refined analysis of hierarchical complexity through different degree strengths in the network. We split each network into four tiers and then eight tiers based on maximum degree magnitudes, where each tier comprised a rounded 25% (12.5% for 8 tiers) of degrees, and so that each tier in the 4-tier split comprised of two tiers in the 8-tier split. The first tier comprised nodes in the top 25% (12.5%) of degrees in the network, the second tier comprised of nodes with the next 25% (12.5%) of largest degrees, and so on. This was implemented on the human brain structural connectomes alongside the randomized structural connectomes to investigate if there were any tiers that were particularly responsible for the differences in hierarchical complexity found. Note that, due to the differences in structural connectomes between subjects, these tiers are not the same for each subject. Making the tiers the same across subjects would obfuscate results as nodes of a certain degree in one network may be regarded as hub nodes, but not so in another network. After this, we computed which regions were consistently—in over two thirds of participants— within one of the tiers to understand the relationship between the cognitive/physiological function of the region and its hierarchical complexity.

### Neighbourhood degree variance

Next, to investigate the diversity of connectivity patterns in the brain, we first isolated notable ROIs by computing the variance of degrees in each region’s neighbourhood. That is, we computed

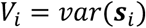

for each tiered region, *i*, and then averaged over participants. We also did this for the random configuration models for comparison. Of these computations, we marked ROIs which were over one standard deviation from the mean within each tier for further analysis. These were targeted as regions with particularly notable structural behaviour. For these we assessed under- and over-representation of tiers. This was done for individual ROIs as follows. The number of neighbouring nodes of the ROI, 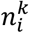, within a given tier, *k*, was noted. The fraction of nodes coming from a single tier was then taken as 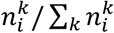, which we called the observed fraction. At the same time, the expected fraction was taken using the number of nodes in each tier within the whole network, *N*_*k*_, as a fraction of the total number of tiered nodes, *N*_*k*_/ ∑_*k*_ *N*_*k*_.

### Statistical Analysis

Population *t*-tests were carried out to assess the significance of the differences of distributions of network index values between adult structural connectomes and random null models as well as between pairs of random null models. The effect sizes were also computed with Cohen’s *d*.

## Results

### Analysis of global binary topology

We first tested the hypothesis that the adult human structural connectome is hierarchically complex by a comparison with relevant null models. We then employed a comparative analysis with a number of well substantiated network indices to understand what new information was provided by hierarchical complexity. The results of hierarchical complexity, *R*, of the binary human structural connectome, alongside more standard measures of network topology— heterogeneity (*V*), assortativity (*r*) and segregation (*γ*)— are plotted against network density (*P*) in Figure 3. The hierarchical complexity of the human structural connectome data is notably larger than the three randomised null models, Figure 3(a). Values of *R* for human structural connectomes have a mean and standard deviation of 0.224 ± 0.055 whereas randomly reconfiguring edges drops *R* by almost half to 0.137 ± 0.030 (effect size of 1.400 with respect to structural connectome values). Additionally, much lower values of *R* are obtained by RGGs (0.087 ± 0.032 with an effect size of 1.670) and random graphs (0.013 ± 0.003 with an effect size of 1.870). The effect size between RGGs and configuration models was 1.261. All of these comparisons drew *p* values of statistical *t*-tests less than 0.0001.

**Figure 3.**
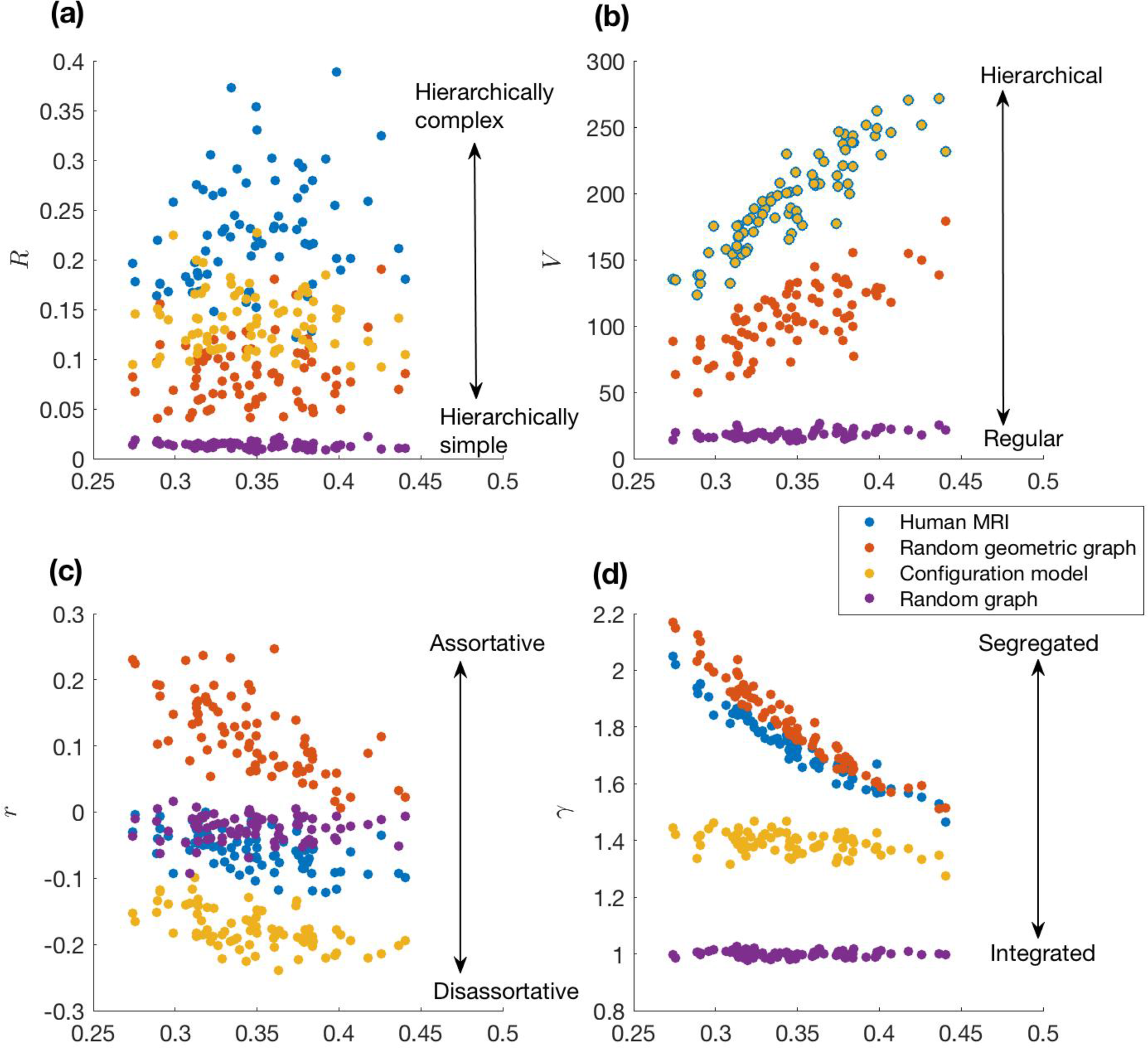
Hierarchical characteristics of the human structural connectome compared to relevant randomised graphs (a-b). Included are the assortativity (c) and random graph normalised clustering coefficient (d) for comparison. While the other characteristics cannot separate all the different network types, hierarchical complexity displays a scale ranging from hierarchically simple Erdos-Renyi (E-R) random networks through random geometric graphs (RGGs), then random networks with the same degree distributions as human MRI networks, and finally to the most hierarchically complex human MRI networks.

Correlations between index values achieved by structural connectomes were computed, including additional computations of characteristic path length and rich-club coefficients. These data show that hierarchical complexity was the least overall correlated index, see supplementary material Section II. On the other hand, values for clustering coefficient, degree variance, characteristic path length and mean rich-club coefficient were all highly correlated indicating that they point to the same topological phenomenon of these networks. Furthermore, index values were assessed for associations with age and sex. Findings indicated that the hierarchical complexity of structural connectomes was independent of these factors, whereas the correlated indices produced a significant effect. This indicated that degree variance and clustering coefficient, in particular, were both higher in older people and in women.

Each null model used takes up a distinct region of the hierarchical complexity spectrum whereas overlaps are present between the MRI data and one of the null models in all of the other spectrums analysed, as shown by the mean ± standard deviation of network measure values presented in Table 1. Conceptual ranges for each network measure are illustrated to the right of the plots in Figure 3. The human structural connectome is the most hierarchically complex of all the models but has the same amount of hierarchical structure (degree variance, Figure 3(b)) as the configuration models since the degree distribution is fixed. As for random graphs, human structural connectomes are neither assortative nor disassortative indicating that nodes of a given degree are neither likely nor unlikely to be connected to nodes of a self-similar degree, Figure 3(c). Finally, as for RGGs, human structural connectomes have similar levels of high segregation, indicating that nodes tend to cluster together in the connections they make in a similar manner to that in distance-based networks, Figure 3(d).

**Table 1.** Mean ± standard deviaton of network measures of brain connectomes and random graph models.

| Table 1. Mean ± standard deviation of network measures of brain connectomes and random graph models. | | | | |
| --- | --- | --- | --- | --- |
| Table 1 | Brain MRI | Randomised MRI | RGG | Random Graph |
| $R$ | $0.224 \pm 0.055$ | $0.137 \pm 0.030$ | $0.087 \pm 0.032$ | $0.013 \pm 0.003$ |
| $V$ | <u>$195.252 \pm 36.233$</u> | <u>$195.252 \pm 36.233$</u> | $106.166 \pm 25.009$ | $18.578 \pm 2.944$ |
| $r$ | <u>$-0.057 \pm 0.029$</u> | $-0.176 \pm 0.029$ | $0.116 \pm 0.058$ | <u>$-0.025 \pm 0.020$</u> |
| $C$ | <u>$0.600 \pm 0.026$</u> | $0.483 \pm 0.048$ | <u>$0.623 \pm 0.019$</u> | $0.347 \pm 0.038$ |

*Note. $R$ : hierarchical complexity, $V$ : degree variance, $r$ : assortativity, $C$ : clustering coefficient. The underlined values in each row indicate cases where standard deviations overlap with each other's means.*

Figure 4 provides an illustration of why the human brain structural connectome has such high hierarchical complexity. In this instance, for 31-degree nodes, the participant’s structural connectome (bottom left) has three nodes with widely varied neighbourhood degree sequences. On the other hand, the random null models have much more homogeneous neighbourhood degree sequences.

**Figure 4.**
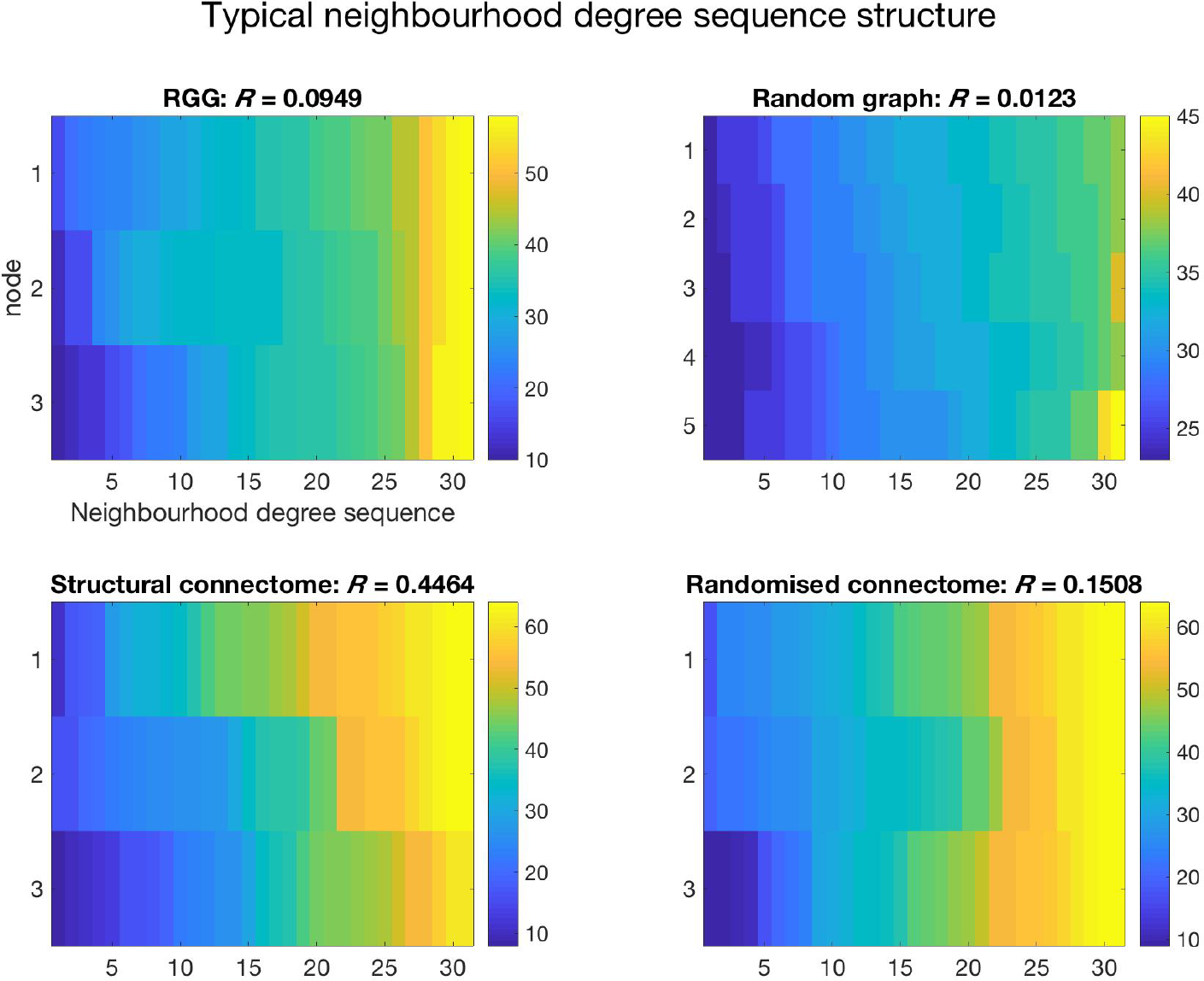
An example of neighbourhood degree sequences of nodes of degree 31 for the structural connectome of a single subject (bottom left) compared to node and edge matched random models. For this subject, the randomized connectome and the RGG there are three nodes of degree 31 in the network whereas for the random graph there are five. Note how each degree sequence in the structural connectome is distinct, whereas degree sequences are far more similar in the random models.

### Analysis of hierarchy tiers

We then examined which nodes in the hierarchy are contributing to greater complexity. To do this we split the nodes up into a number of tiers based on their degrees and looked at the effect size of hierarchical complexity within tiers between structural connectomes and edge-randomised connectomes. Table 2 displays these results for 4-tier and 8-tier strategies, with the individual data points plotted per subject in Figure 5. An analysis of regional (ROI) consistency within the tiers can be found in the supplementary material, Section III, alongside the reported mean and standard deviations of the mean degree within each tier.

**Table 2.** Effect size (Cohen’s *d*) of hierarchical complexity, *R*, within tiers between structural and randomised connectomes.

|  |  |  |  |  |  |  |  |  |
| --- | --- | --- | --- | --- | --- | --- | --- | --- |
| # of Tiers | Tier 4b | Tier 4(t) | Tier 3b | Tier 3(t) | Tier 2b | Tier 2(t) | Tier 1b | Tier 1(t) |
| 4-Tier | - | 0.754** | - | 1.320** | - | 1.105** | - | 0.491 |
| 8-Tier | 0.489 | 0.702** | 1.043** | 1.110** | 0.821** | 0.501* | 0.3432 | 0.120 |

Note. \* denotes population *t*-test with $p < 0.01$ , \*\* denotes population *t*-test with $p < 0.0001$

**Figure 5.**
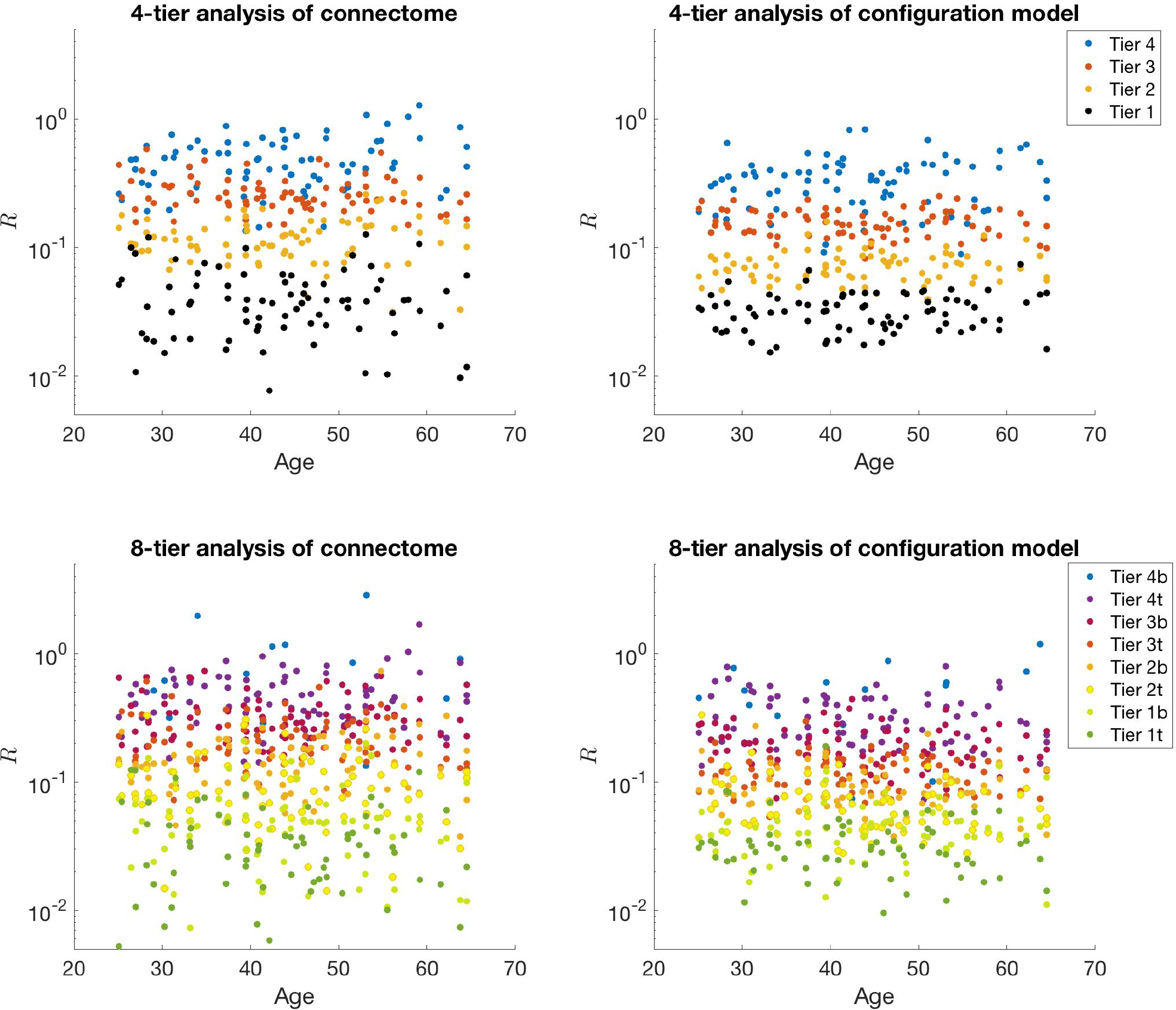
Analysis of hierarchical tiers contributing to the high hierarchical complexity in the human structural connectome, left, compared to their random configuration models, right, for 79 individuals. Given *T* tiers, Tier 1 comprises the 100/*T* % most highly connected nodes whereas the final tier is the 100/*T* % of nodes with the smallest degrees.

The results show that hub nodes (Tier 1(t)) and peripheral nodes (Tier 4(b)) are contributing less to the greater complexity exhibited in the human brain connectome than middle tiers. In fact, this is particularly true of hub nodes, with lowest effect sizes notable in Tier 1 and Tier 1t in respective 4- and 8-tier strategies. Indeed, the additional tiers in the 8-tier analysis— splitting each 4-tier tier into a top (t) and bottom (b) tier— shows that there is no significant difference between configuration model and human brain data in Tier 1t, corresponding to the top half of Tier 1 in the 4-tier analysis (population *t*-test *p* = 0.492, *t* = 0.690). However, the bottom half of Tier 1 in the human connectome has an effect size over two and a half times greater and does show a slight significant difference to the configuration model (*p* = 0.043, *t* = 2.040). The same pattern repeats itself in the analysis of the last tier where the extremity shows least difference. The difference between human brain and randomised data for the Tier 4b nodes in the 8-tier analysis had *p* = 0.179 (*t* = 1.378), whereas the difference found in Tier 4t had *p* < 0.0001 (*t* = 4.611).

The ROIs relating to the four tiers—those which are in a given tier in more than two-thirds of participants—are as in Table 2. Such consistency was found for over 70% of brain ROIs (61 of 83).

**Table 2.**
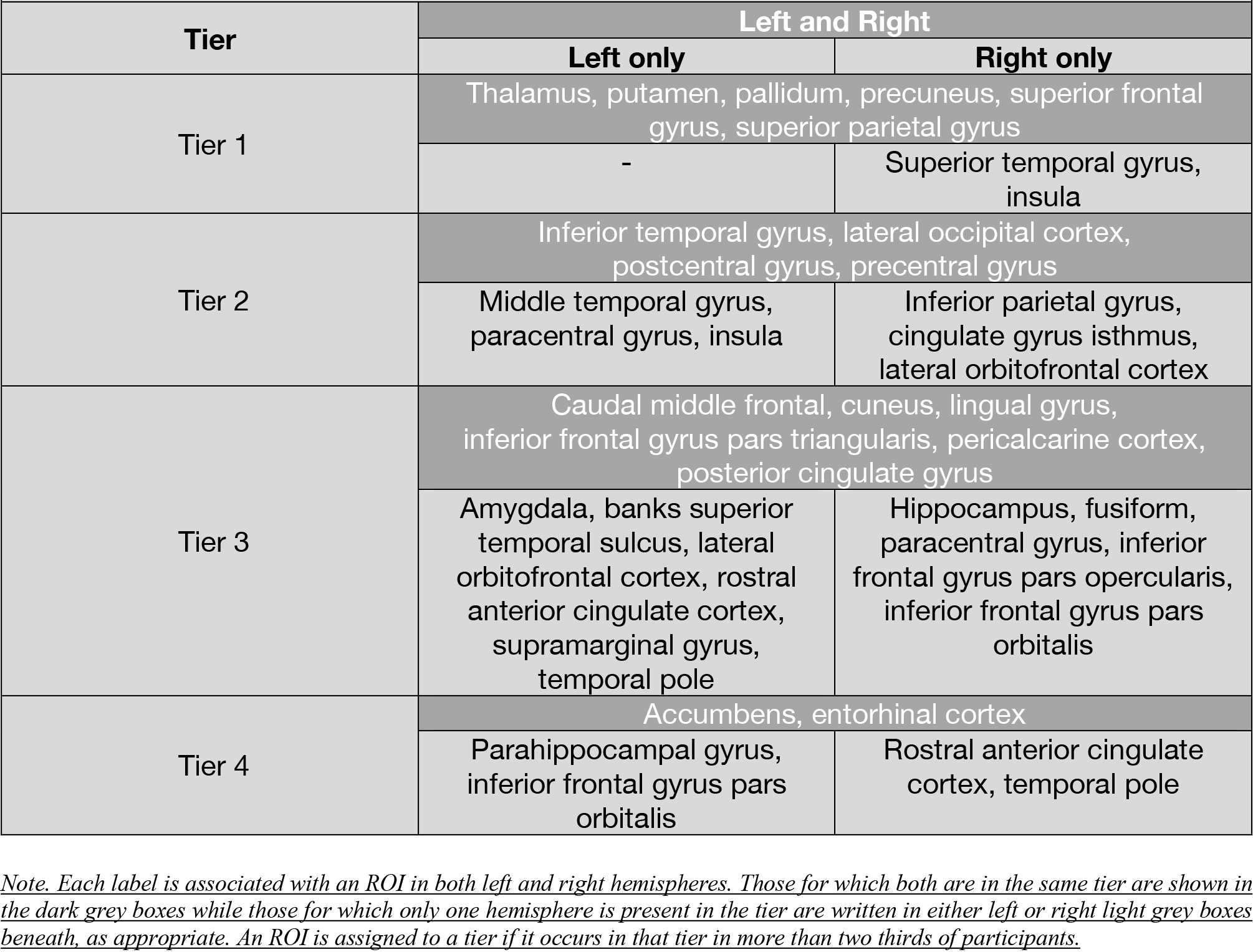
Classification of brain ROIs into hierarchical tiers.

| Table 2. Classification of brain ROIs into hierarchical tiers. |  |  |
| --- | --- | --- |
| Tier | Left and Right |  |
|  | Left only | Right only |
| Tier 1 | Thalamus, putamen, pallidum, precuneus, superior frontal gyrus, superior parietal gyrus |  |
|  | - | Superior temporal gyrus, insula |
| Tier 2 | Inferior temporal gyrus, lateral occipital cortex, postcentral gyrus, precentral gyrus |  |
|  | Middle temporal gyrus, paracentral gyrus, insula | Inferior parietal gyrus, cingulate gyrus isthmus, lateral orbitofrontal cortex |
| Tier 3 | Caudal middle frontal, cuneus, lingual gyrus, inferior frontal gyrus pars triangularis, pericalcarine cortex, posterior cingulate gyrus |  |
|  | Amygdala, banks superior temporal sulcus, lateral orbitofrontal cortex, rostral anterior cingulate cortex, supramarginal gyrus, temporal pole | Hippocampus, fusiform, paracentral gyrus, inferior frontal gyrus pars opercularis, inferior frontal gyrus pars orbitalis |
| Tier 4 | Accumbens, entorhinal cortex |  |
|  | Parahippocampal gyrus, inferior frontal gyrus pars orbitalis | Rostral anterior cingulate cortex, temporal pole |

*Note. Each label is associated with an ROI in both left and right hemispheres. Those for which both are in the same tier are shown in the dark grey boxes while those for which only one hemisphere is present in the tier are written in either left or right light grey boxes beneath, as appropriate. An ROI is assigned to a tier if it occurs in that tier in more than two thirds of participants.*

These have been mapped to the MRI image in Figure 6 with different colours representing the different tiers. The same computations were applied to the 8-tier split, however very little regional consistency was found within tiers (only 20%-17 of 83- ROIs could be classified), suggesting that the 4-tier strategy provided the right size for such analysis.

**Figure 6.**
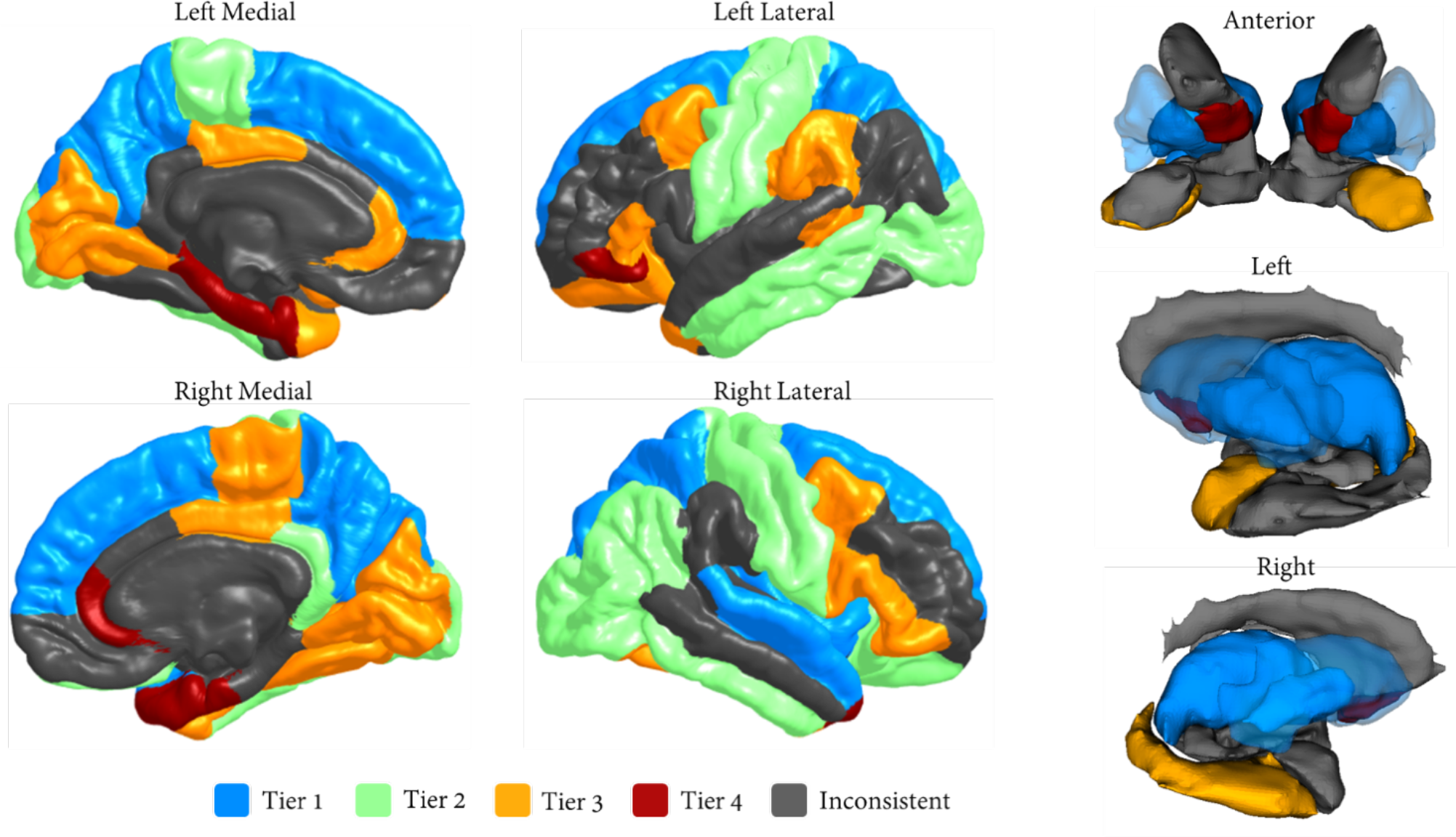
Cortical (left) and subcortical (right) mapping of hierarchical tiers. Grey denotes areas that did not appear in any tier in more than two thirds of participants. Putamen is opaque to enable visualisation of the pallidum.

### Analysis of ROI tier connectivity profiles

Here, we performed an analysis using neighbourhood degree variances to determine particularly notable connectivity patterns within ROIs. The relative neighbourhood degree variances for each ROI (value for brain ROI – value for configuration model ‘ROI’) were plotted onto a cortical surface map, Figure 7. The absolute values for each ROI and their configuration model counterparts can be found in Section IV of the supplementary material. Configuration models had values most likely independent of the average degree of the ROI and did not vary by much. On the other hand, brain ROIs had a very large range of neighbourhood degree variances as can be seen by the red and blue ROIs in Fig 7. Notably, the ROIs in Table 3 were found to be outside one standard deviation of the mean within its tier except in Tier 4 where two clear clusters of large variance and expected variance were found. Two very clear observations could be made here. Firstly, tiers 1 and 4 (the least overall complex) showed strong hemispheric symmetry within these categorisations. Secondly, the ROIs in tiers 2 and 3 (the most overall complex) were almost entirely from the right hemisphere, indicating that right hemisphere ROIs are more diversely connected than left hemisphere ROIs.

**Figure 7.**
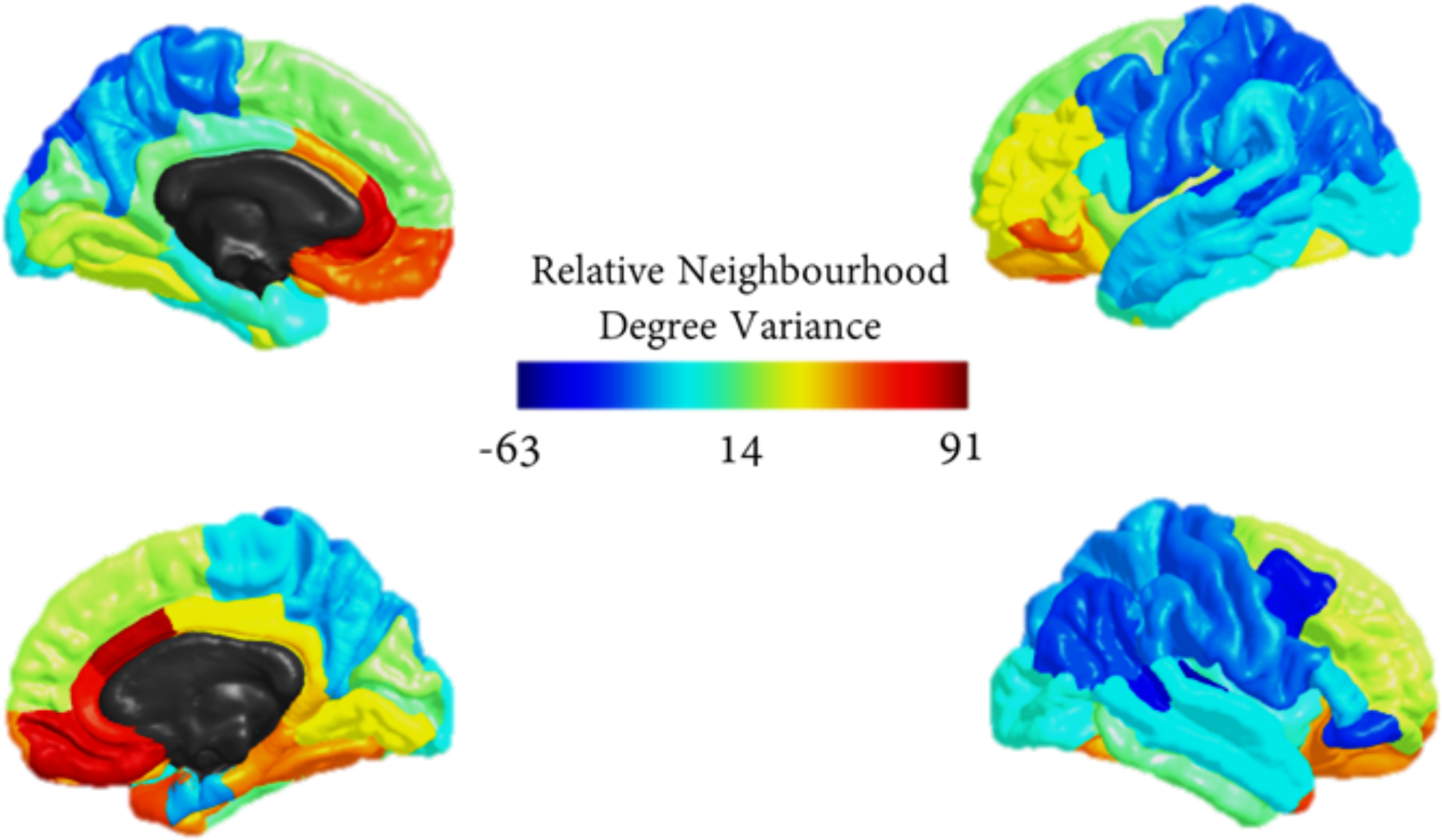
Average neighbourhood degree variance over participants for individual ROIs— relative to values obtained for configuration models— plotted as intensities on a cortical map.

**Table 3.**
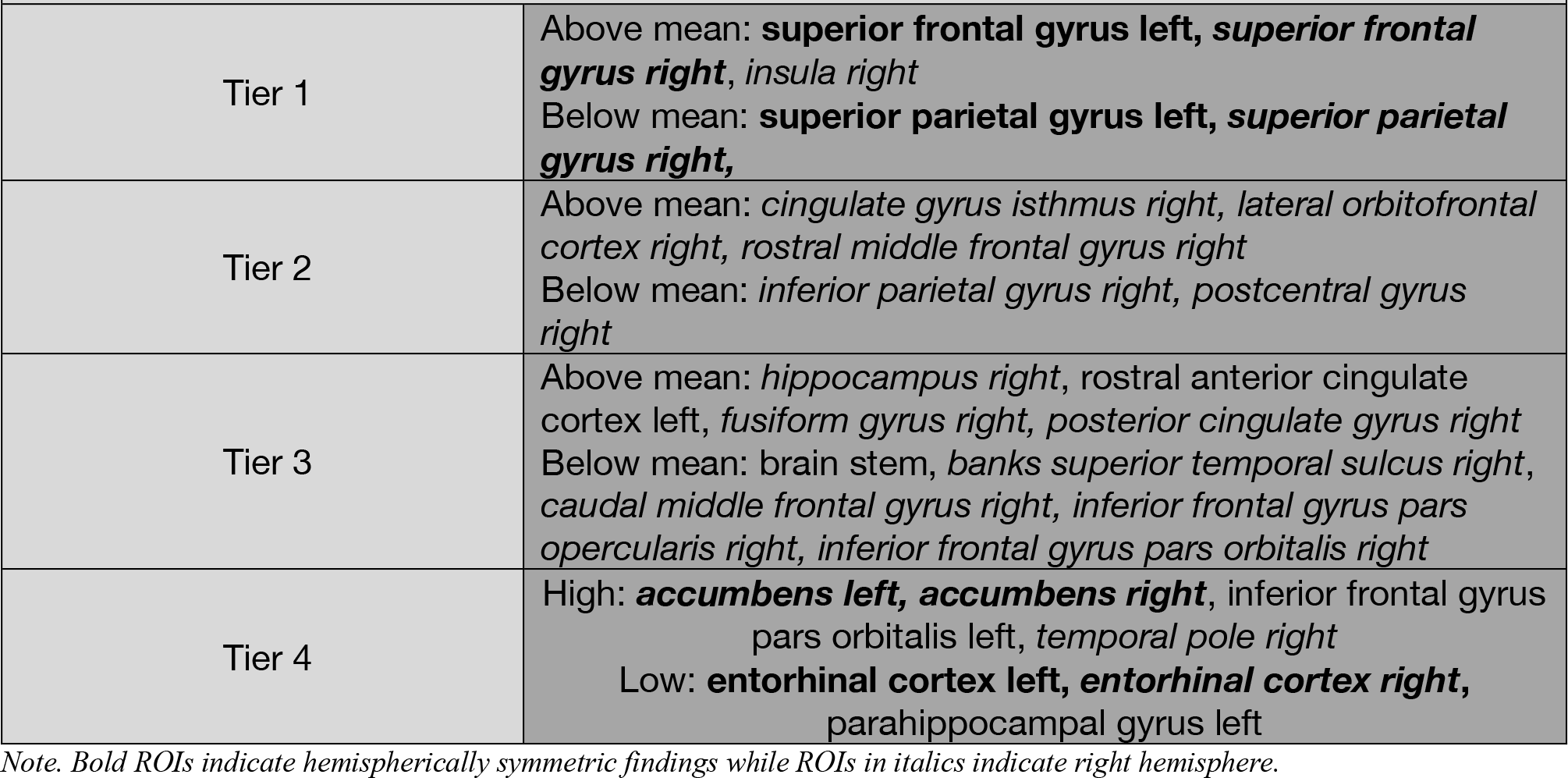
ROIs with neighbourhood degree variance outwith one standard deviation of the tier mean.

We then looked more closely at tier connectivity profiles of notable ROIs to substantiate possible biological reasons for their structural configurations. Average fractions of neighbourhoods within each tier were computed for the symmetric findings in Tiers 1 and 4— superior frontal gyrus, superior parietal gyrus, accumbens and entorhinal cortex— alongside ROIs with particularly high variance—cingulate gyrus isthmus right, lateral orbitofrontal cortex right, hippocampus right, rostral anterior cingulate cortex left, fusiform gyrus right—and low variance—brain stem, caudal middle frontal gyrus right—shown in Fig 8. Table 6 in the supplementary material shows the effect sizes between the observed and expected distributions of neighbourhoods amongst tiers of these ROIs. With few exceptions, the observed distributions were significantly different from the expected distributions. The brain stem showed the largest representation from Tier 1 nodes (Cohen’s *d* = 1.74) whereas the left and right entorhinal cortices were the only ROIs studied which had under-representations of Tier 1 nodes (non-significant *d* = −0.32 for left and significant *d* = −0.66 for right). The greatest representation of Tier 2 nodes occurred in the superior parietal gyrus (*d* = 1.49 for left and *d* = 1.44 for right) whereas only the left rostral anterior cingulate cortex and right entorhinal cortex showed significant under-representations (*d* = −0.71 and *d* = −0.67, respectively). The only over-representations of Tier 3 nodes occurred in the left and right entorhinal cortex (*d* = 0.62 and *d* = 0.43, respectively), whereas the greatest under-representations were found in the brain stem (*d* = −1.78), right fusiform gyrus (*d* = −1.64) and right hippocampus (*d* = −1.58). Finally, the only significant over-representations of Tier 4 nodes occurred in the left rostral anterior cingulate (*d* = 0.97) and right entorhinal cortex (*d* = 0.67), whereas greatest under-representations were found in the brain stem (*d* = −1.90), right caudal middle frontal gyrus (*d* = −1.89) and both left and right superior parietal gyrus (*d* = −1.83 and *d* = −1.80, respectively).

**Figure 8.**
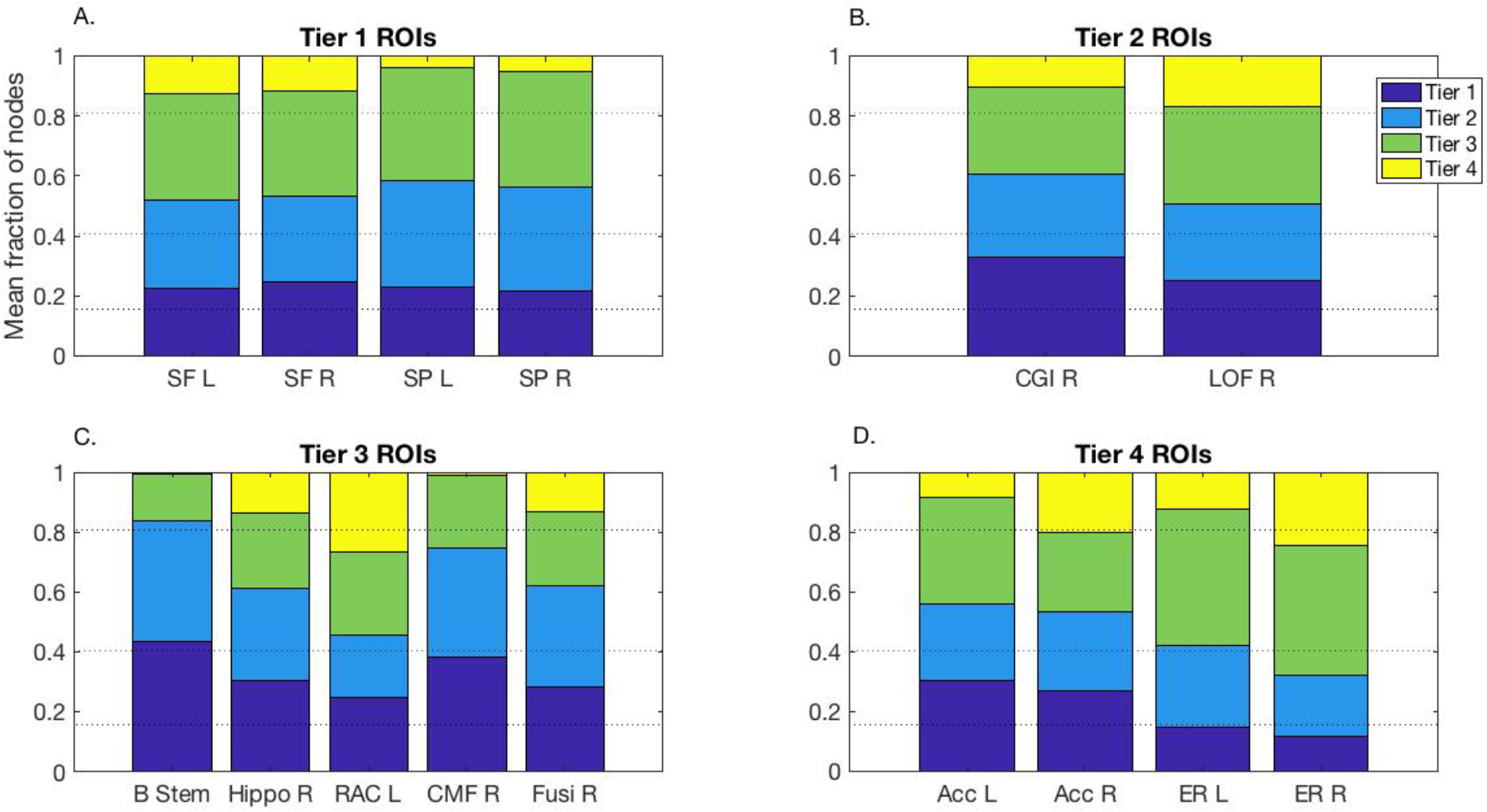
Fraction of neighbouring nodes from tiers in selected ROIs. Dashed lines indicate the fractions of nodes in each Tier, representing the expected fractions; L/R- Left/Right; SF- Superior Frontal gyrus; SP - Superior Parietal gyrus; CGI- Cingulate Gyrus Isthmus; LOF- Lateral OrbitoFrontal gyrus; B Stem- Brain Stem; Hippo- Hippocampus; RAC- Rostral Anterior Cingulate cortex; CMF- Caudal Middle Frontal; Fusi- Fusiform; Acc- Accumbens; ER- Entorhinal cortex.

## Discussion

We confirm the hypothesis that human brain structural connectomes created from structural and diffusion MRI data are more hierarchically complex than random null network models. It is interesting to note that randomising edges in networks with identical degree distributions to those obtained from brain MRI data — thus fixing degree variance— provides networks with a dramatically decreased hierarchical complexity. This indicates that the dissimilarity of connections made by network nodes with the same centrality cannot be explained by greater variability of network degrees present and shows a prominent presence in the brain structure of dissimilarity between the nodes residing at the same hierarchical level. This suggests that the organisational complexity in the human brain is more heterogeneous than that produced at random and that heterogeneity in the connectivity patterns of hierarchically equivalent nodes could itself yet prove a single coherent explanation for the complexity of brain structure.

Indeed, EEG functional connectivity was found to be more complex than a variety of ordered systems as well as Erdos-Renyi random networks. However, hierarchical complexity depends on degree distributions and it had yet to be shown whether or not brain networks were more hierarchically complex than random networks with the same degree distributions as brain networks. The data presented here is thus the strongest evidence yet to support the hierarchical complexity paradigm as the key to distinguishing real-world complexity from the more predictable patterns of ordered and random systems. To test this theory further, it would be of significant interest to implement similar analyses on other brain signals/imaging such as in a cortical stimulation study, and in other modalities such as magnetoencephalography and functional MRI.

From a neuroanatomical perspective, the tiers from the 4-tier categorisation exhibited a degree of anatomical plausibility. Tier 1 (highest degree but lowest contribution to hierarchical complexity) comprised lateral frontal, parietal and lateral temporal regions along with selective subcortical structures. This corresponds well with the current (macro)neurobiological account of intelligence differences (the Parieto-Frontal Integration or P-FIT theory^30–32^) and resembles previous work which identifies hub nodes of the human brain connectome^16^. Tier 2, on the other hand, consists mainly of occipital and sensorimotor cortex involved in lower order sensory processing. Interestingly, Tier 3 is then comprised of mainly heteromodal integrative regions which may represent a transitional stage in information processing between higher order cognitive (Tier 1) and lower order sensory processing (Tier 3)^33^. Functional categorisation of Tier 4 ROIs is less clear-cut, but the nucleus accumbens, entorhinal cortex and anterior cingulate are all ostensible components of the hippocampal-diencephalic-cingulate network involved in memory and emotion^34^.

Crucially, our analyses suggest that hierarchical complexity is not driven by hub nodes (Tier 1), but rather by nodes particularly in Tier 3 (mainly heteromodal integrative regions) and to a lesser but still significant extent in Tier 2 (more basic sensorimotor and visual-semantic areas). Given that Tier 3 consists of ROIs involved in collecting/integrating information from both ends of the processing spectrum (higher order cognitive and more basic sensory), the great diversity of cross-tier connectivity patterns revealed by its large hierarchical complexity stands to reason. Indeed, it suggests a rich diversity of roles played by these integrative regions, necessitating connections across all tiers. The fact that hub nodes (Tier 1) are not found to be substantial contributors to the hierarchical complexity of human structural connectomes indicates that hub nodes may take on a disproportionate amount of focus in brain network studies^35^. These findings raise interesting prospects to see how such diversity (or lack thereof) of cross-tier connectivity patterns are affected in pathology and disease. Particularly appropriate would be to apply these methods to diseases known to affect different steps of multimodal functional integration (cognition, sensorimotor, or the integration of these processes). It is important to keep in mind, however that not all participants have exactly the same tier structure, thus any generalisation of these results should be made with due caution.

From neighbourhood variance analyses, it is interesting to note the high diversity found within right hemisphere ROIs, indicating asymmetric differences. It is also important to remember that all subjects were right-handed. Functional inter-hemispheric differences and differential hemisphere specialisation are well known. Whilst the left hemisphere has been found to be dominant for speech, the right hemisphere is known to play a major role in many non-verbal cognitive functions, and particularly in the perception of spatial relations^36^. Sex-related inter-hemispheric differences have also been reported^37,38^ although a large study presented the idea of a “brain mosaic” of features, some more common in females compared with males and vice versa, after analysing MRI, personality traits, attitudes, interests, and behavioural data from more than 5,500 individuals^38^. In our own experience, for example, hippocampal shape deformations in relation to cognitive functioning exhibit also a high degree of asymmetry^39^. Hierarchical complexity may well provide a link to understand the mechanisms and targeting behind such asymmetric properties.

We found that certain ROIs were well integrated across the hierarchy, whereas other ROIs were more selective and connected to a more limited hierarchical range. For example, the low neighbourhood degree variance of the brain stem was found to be caused by a large under-representation of Tier 4 and 3 (lowest degree) nodes and a large over-representation of Tier 1 hub nodes, Fig 8C. The brain stem itself being a Tier 3 node tells us that it communicates ‘upwards’, i.e. more exclusively to those ROIs in higher tiers (cognition and sensory processing). The same was true for the caudal middle frontal gyrus of the right hemisphere. On the other hand, the high variance associated with the rostral anterior cingulate cortex of the left hemisphere was caused by over-representations of the top and bottom tiers, which, from our descriptions of tiers, agrees with the understanding of its integrative role in cognition and emotion and the known heterogeneity of structural connectivity in this portion of the cingulate gyrus^41,42^. The entorhinal cortex displayed neighbourhood degree variance similar to those of the configuration models and, indeed, the tiers were all fairly well represented for the entorhinal cortex, indicating this region was very well integrated throughout the brain. Particularly, this ROI being the only highlighted region which had over-representations of Tier 3 nodes indicates a key role in cognitive and sensory integration which lines up with the knowledge that superficial layers of the entorhinal cortex receive both multimodal and unimodal sensory inputs while also projecting to widespread cortical and subcortical loci^43^. Also, the large representations of Tier 2 nodes in the superior parietal gyrus aligns with its known predominant function in sensorimotor stumuli^44^.

Hierarchical complexity can also help deepen our understanding of other topological findings in the connectomes. For example it provides an explanation for the un-assortative nature of brain structural connectomes^45^, (see Figure 3(c)). The degree of nodes in a given node’s neighbourhood do not maintain a self-similarity to the degree of the given node, because nodes take up a wide array of different neighbourhood connectivity patterns encompassing the heterogeneity of degrees in the whole network. Including results of the high values of *γ*—propensity of neighbouring nodes to share other neighbours— indicates that i) nodes which are connected together tend to connect to the same other nodes (high *γ*), ii) these nodes do not have similar degrees, (*r* ≈ 0) and iii) nodes of the same degree do not have similar distributions of neighbouring degrees (high *R*). All of these aspects are somehow integrated into the brain connectivity structure to create this rich and diverse topology.

It is also interesting to note a striking overlap in segregation between the human structural connectome and RGGs (see Figure 3(d)). The strength of this overlap, together with the lack of hierarchical structure present in RGGs, suggests that geometric sensibilities of node clustering is extended also to integrative connections, where two connections spanning the connectome within a geometrical locality tend to span to the same nodes in the other locality. This agrees with the homophily principle described in a recent connectome simulation study^9^. Note that the results here significantly differ from another study where segregation in RGGs was found to be larger than the connectome^7^, although it must be noted that the network size (*n* = 998) was much larger and density (*P* = 2.7%) more sparse than the current study and the space used to develop the models was rectangular rather than cubic as adopted here. It should also be noted that sparsity is not a desirable feature for analyzing hierarchical complex networks^11^.

This is particularly interesting in the context of pathology. It is not yet clear which measure (or measures) can explain functional outcomes from pathological features (i.e. lesions, mineral accumulations, tissue loss, etc.), an understanding of which is required to help solve what has been termed the clinico-radiological paradox^46^. Evidence shows that there are specific white matter pathways that have greater impact on clinical and functional outcome regardless of the lesion size^47^ whilst other tracts offer routes for functional reorganisation^48–50^. Future studies applying the hierarchical complexity measure to health and disease may help to uncover subtler but still significant differences in brain network topology that will add to our understanding of this topic. For example, we generally expect that brain degradation (whether from ageing or disease) will display structural connectivity patterns more similar to the edge-randomised networks. The supplementary results showing that hierarchical complexity is independent of age and sex and is not highly correlated with other indices, indicate that hierarchical complexity is a unique factor of topology which may be maintained in the face of other topological variables. We conjecture that this thus underlines structural characteristics of fundamental importance for the emergence of complex integrated brain function. In the future we aim to test this hypothesis by looking for associations of hierarchical complexity with intelligence and pathology.

The evidence here adds to previous results of hierarchical complexity found in EEG functional connectivity^10,11^, revealing a topological agreement in complexity between structure and function— both being more hierarchically complex than a variety of pertinent models. Future studies on the relationship between structural and functional MRI with respect to this complexity paradigm will help to better understand how function relates to structure and whether the structural complexity found here supports complex functional principles. Additionally, the aggregated tissue of individual ROIs here are abstracted as network nodes, however it would be of high relevance to look at whether hierarchical complexity is a self-similar property of brain networks by considering different scales of brain networks^51^.

One limitation of the study is that we have not shown invariance of connectome hierarchical complexity to the choice of parcellation scheme. Different atlases and methods for producing connections do exist, but the resulting networks have been found to broadly share topological characteristics (e.g. small-world and degree distributions), even if the exact values of indices between different schemes are statistically different^52^. In addition, we previously demonstrated that different sizes of EEG functional networks share the characteristic of hierarchical complexity^10^, suggesting that results may not significantly differ when using other parcellation schemes in structural MRI, notwithstanding general effects of parcellation granularity on tractography results^53^. Another possible limitation is that the data were collected at 1.5 rather than 3T (or above). Higher field strengths have the potential to provide better tractography and parcellation information due to increased signal to noise ratio. However, higher field strength also has a greater potential for artefacts and does not necessarily result in better diagnostic accuracy^54^. Furthermore, consistency has been found in connectivity profiles across field strength^55^. Additionally, 1.5T scanning is still widely used in both clinical and large prospective cohort studies^40,56–58^.

## Conclusion

The adult human structural connectome was found to be hierarchically complex with highly heterogeneous connectivity patterns occurring across hierarchically equivalent nodes. This was established in comparison to three very different random models—Erdös-Rényi random graphs, RGGs and edge-randomised connectomes. Hierarchical complexity was shown to divide the different models into a coherent range of topology with the human structural connectome at the top, while other standard topological concepts of segregation, assortativity and heterogeneity failed to adequately separate the models. These data suggest that diversity of connectivity patterns of hierarchically equivalent nodes could itself provide a cohesive rule for generative processes of brain structure. Moreover, this may explain the difficulty in establishing accurate generative models which account for all aspects of brain connectome topology using more predictable patterns. Hierarchical complexity was most apparent in Tier 2 and 3 nodes, constituting brain regions involved in sensorimotor, attentional and linguistic-semantic function, whereas tiers 1 (hub nodes related to general intelligence) and 4 contributed much less to the complexity. Tiers 1 to 3 mapped to the different steps of the proposed functional connectivity framework for the integration of cognitive and sensory processing. From this, the most hierarchically complex tier contained the ROIs involved in the integration of cognitive and sensory inputs. These results were supported by specific neighbourhood analyses by tiers which found structural configurations of neighbourhoods which aligned function of specific ROIs with this integrative processing framework. This study provides a platform from which to explore hierarchical complexity of the human structural connectome in cognition, health and disease.

## Supporting information

Supplementary analyses

## Acknowledgements

This work is dedicated to the memory of Prof. John M. Starr. This work was supported by Health Data Research UK (MRC ref Mr/S004122/1), which is funded by the UK Medical Research Council, Engineering and Physical Sciences Research Council, Economic and Social Research Council, National Institute for Health Research (England), Chief Scientist Office of the Scottish Government Health and Social Care Directorates, Health and Social Care Research and Development Division (Welsh Government), Public Health Agency (Northern Ireland), British Heart Foundation and Wellcome. MCVH is funded by the Row Fogo Charitable Trust (Grant BRO-D.FID3668413). Data collection was funded by NIH grant R01 EB004155.

## Conflict of interests

The authors declare no competing interests.

## Author Contributions

KS devised the study, computed the network analyses and results and wrote the manuscript. MB collected the MRI data and generated the structural connectomes and helped write the manuscript. SC provided biological interpretation of the results and helped write the manuscript. MVH helped devise the study, provided expertise of MRI data and reviewed the manuscript. SW helped devise the study and reviewed the manuscript. JE helped interpret the technical results and reviewed the manuscript. CS helped devise the study and reviewed the manuscript.

