## Supplementary analyses for "Hierarchical Complexity of the Adult Human Structural Connectome"

### I. Introduction

This document describes supplementary analyses for our article on the hierarchical complexity of the adult human structural connectome. This covers i) extensions of the analysis to rich-club and small-world characteristics and their relationship to hierarchical complexity, ii) associations of network indices with age and sex of the participants of the study, iii) an analysis of regional consistency within hierarchical tiers across participants and the mean degree of regions within each tier, and iv) additional information for the neighbourhood degree variance.

### II. Extended network analyses and associations with age and sex

Characteristics of small-world[1] and rich-club[2] networks have previously been found in structural connectomes. It is important, therefore, to consider whether there is any relationship between hierarchical complexity and such network characteristics in our study.

The small-world network is characterised by a tendency to having strong interconnected clusters of nodes whilst at the same time having an efficient communication enabled by connections between these clusters[3]. The clustering coefficient, described in the main body of text, helps to quantify clustering, whilst the characteristic path length, defined as

$$L = \frac{\sum_{i,j=1}^{n} d(i,j)}{n(n-1)}$$

where $d(i,j)$ is the shortest number of edges required to construct a path between nodes $i$ and $j$ in the network, helps to quantify efficiency, i.e. how quickly can one get from any given node to any other on average. The small-world coefficient is then defined as the ratio of these values relative to the value obtained for random graphs[4]:

$$\sigma = \frac{C/C_{ran}}{L/L_{ran}}.$$

The rich-club coefficient at degree $k$ is defined as

$$K_k = \frac{1}{n_k(n_k-1)} \sum_{i=1}^{n_k} \sum_{j=1}^{n_k} a_{ij}^k,$$

where $A^k$ is the reduced adjacency matrix of $A$ including only those nodes of degree at least $k$ in $A$, with size $n_k$ and entries $a_{ij}^k$. That is, the rich-club coefficient is the density of the subgraph $A^k$ which measures the interconnectedness of the nodes of degree greater than $k$, thus can help tell us the extent to which high degree 'hub' nodes are interconnected[5].

Further to this, to see whether rich-clubs inform on hierarchical complexity, we constructed a network configuration model with a fixed rich-club. This was done by combining the connections made by the top 30 degree nodes of the brain networks with randomised configurations making up the remainder of edges, i.e. randomising only those connections between the bottom 53 degree nodes.

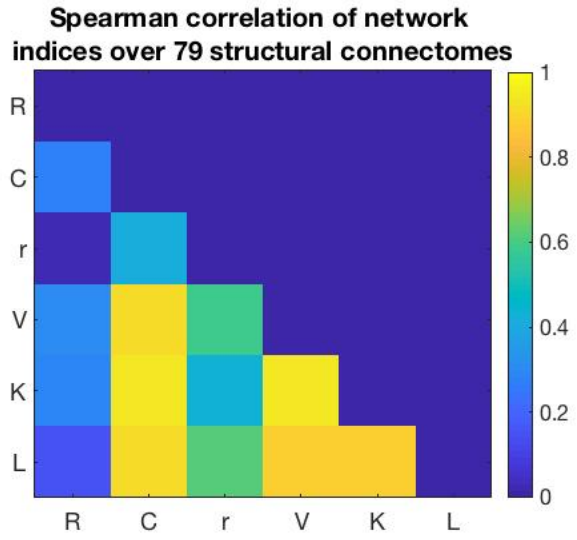


Figure 1. The lower triangle shows absolute value of spearman correlations of different network indices across the structural connectomes. R- hierarchical complexity, C- clustering coefficient, r- assortativity, V- degree variance, K- mean rich-club coefficient, L- characteristic path length. Diagonal and upper triangle are set to zero.

For the statistical analysis, first of all we computed Spearman correlations between the index values obtained for each of the 79 brain structural connectomes for hierarchical complexity, and the 5 other important indices including characteristic path length, $L$, and the mean rich-club coefficient over all $k$, $K$, Fig 1. Two things are immediately notable. Firstly, hierarchical complexity is the least correlated index amongst these. This strongly suggests that hierarchical complexity is a largely independent index which is telling us something unique about human brain structure. Secondly, all of the other indices except from assortativity are highly correlated with each other. This tells us that, contrary to expectations, these network indices, designed to study different characteristics, are all essentially pointing towards a single phenomenon in these connectomes— the heterogeneity of degrees ($V$) in the connectome is highly correlated with the amount of clustering ($C$), the strength of the rich-club ($K$) and the extent of integration found ($L$).

We then computed the mean rich-club coefficients, small-world coefficients and hierarchical complexity for the configuration models and Random Geometric Graphs (RGGs), described in the main document, alongside the rich-club-fixed configuration model described above. The mean rich-club coefficient, indeed, did not deviate significantly between the structural connectomes and the rich-club-fixed model, Fig 2 A. However, hierarchical complexity still significantly decreased even after connections made by the top 30 degree nodes were fixed, Fig 2 C. This tells us that the interactions between low degree nodes is a non-negligible and complex feature of brain networks. Furthermore, surprisingly, small-world coefficients of structural connectomes were found not to be statistically different to those of RGGs. But, again, hierarchical complexity was clearly lower amongst RGGs. Similarities in network topology between RGGs and structural connectomes may be reconciled in light of the known low wiring cost (regions closer together are more likely to be connected together) associated with brain networks[6], while the dramatic difference in hierarchical complexity tells us that the principle of homophily alone, seen as a fundamental property of brain networks[7], does not explain the hierarchical complexity of the structural connectome. Distributions of mean rich-club coefficients and characteristic path lengths among connectomes and models found very high amounts of overlap which were not found to shed more light on the descriptions of topology set forth in Fig 2 of the main document, Fig 2 D & E.

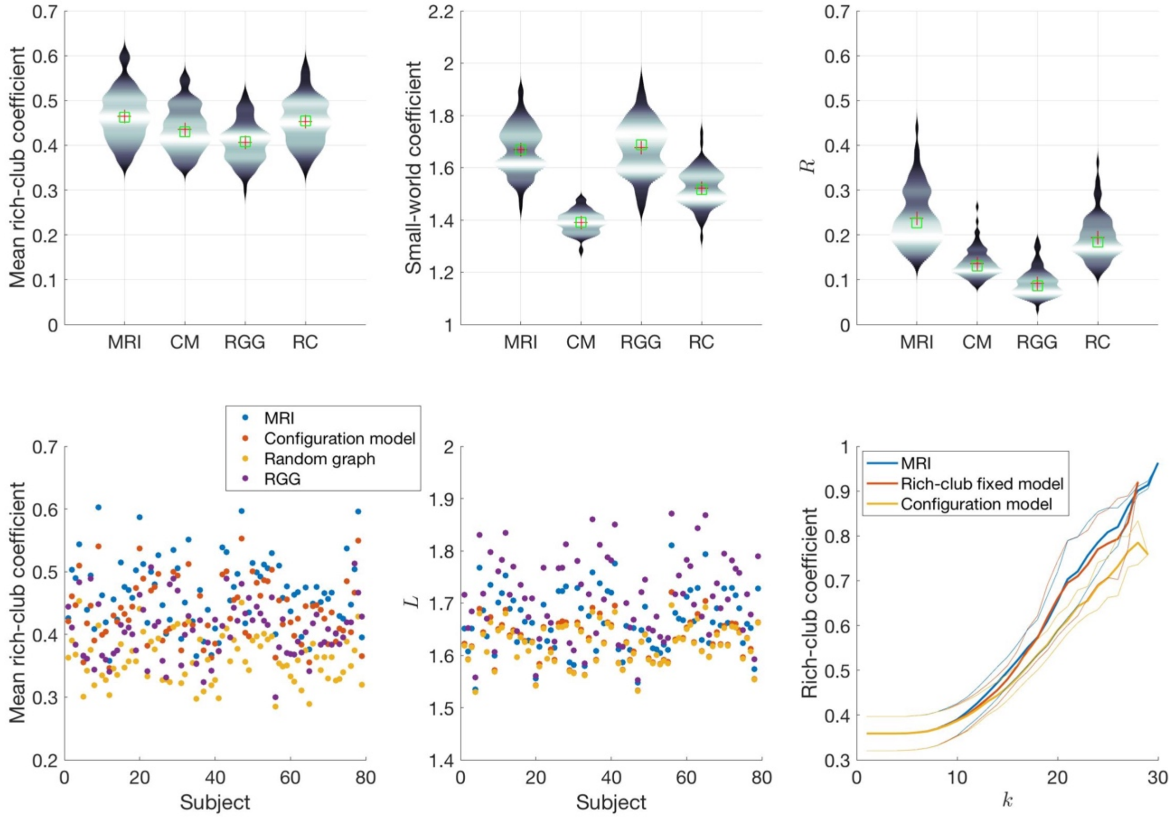


Figure 2. A. Mean rich club coefficient, B. small-world coefficient and C. hierarchical complexity of structural connectomes (MRI), their configuration models (CM), random geometric graphs (RGG) and rich-club fixed configuration models (RC). D. Mean rich-club coefficients and E. characteristic path lengths of structural connectomes (MRI), and different models, showing that these indices are not particularly useful at separating topologies. F. The rich-club coefficient by degree (k) of the rich-club fixed configuration model compared to MRI and regular configuration model.

Next, we sought to compare distributions of index values amongst men and women in different age groups using Mann-Whitney U tests. Figure 3 shows network index values of structural connectomes. Age groups were defined as those under 35 ($n = 21, f{:}m = 10{:}11$) those between 35 and 45 ($n = 23, f{:}m = 12{:}11$), those between 45 and 55 ($n = 23, f{:}m = 11{:}12$) and those over 55 ($n = 12, f{:}m = 5{:}7$). Two-way anova tests were conducted across age (3 degrees of freedom), sex (1 d.f.) and their interaction (3 d.f.) for each index. Results are shown in Table 1. All significant differences passed the false detection rate procedure. Significant differences were found with respect to both age and sex for $C$, $V$ and $L$ and a significant difference was found with respect to age for $K$. No interaction effects were found between age and sex. Clustering and degree variance were higher for older people and for women, while characteristic path length was lower for both and the rich-club was stronger in older people. Of course, all of these indices were found to be highly correlated, so the overlap of results is not surprising. On the other hand, hierarchical complexity and assortativity were not found to differ between men and women nor between age groups. This puts forward the interesting proposition that R and r are properties which are more prioritised within brain structure, as they are roughly the same across age groups and the sexes. They are therefore also potentially very useful to consider for studies in cognitive decline and pathology where invariance to ageing is often highly desirable.

*Table 1. Results (p-values with F statistics in brackets) from 2-way ANOVA tests across age and sex of network indices*

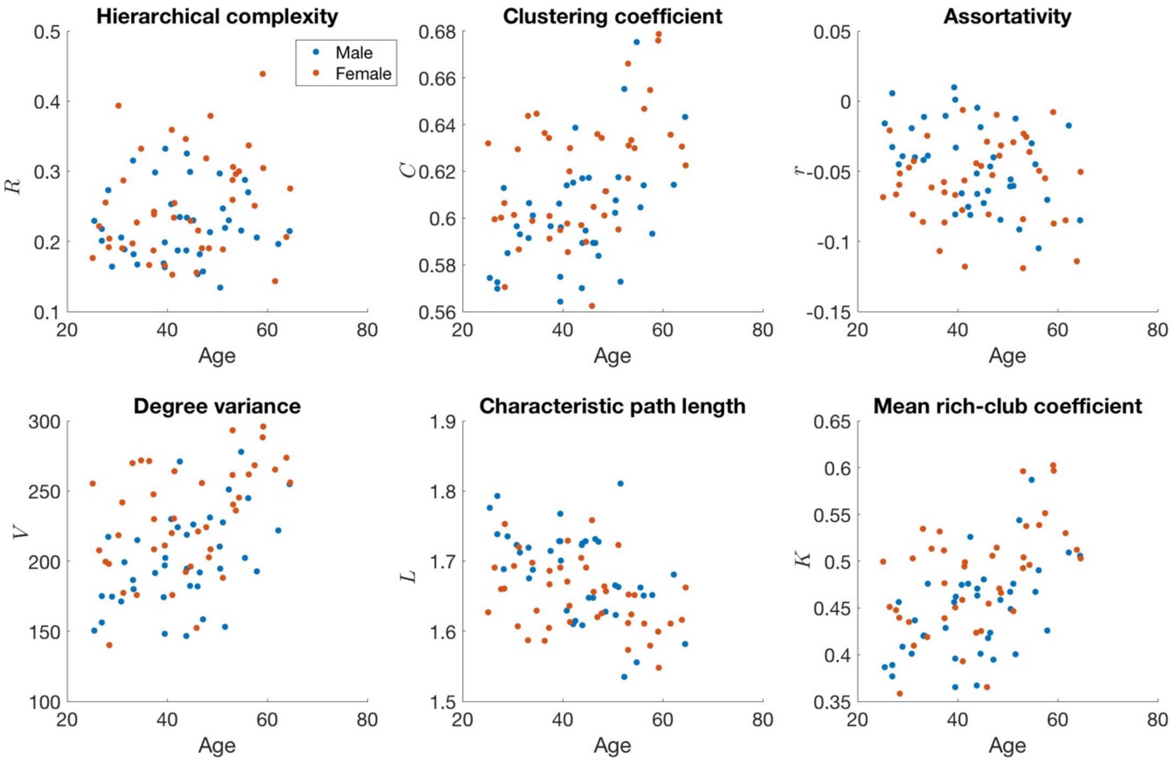


Figure 1. Network indices plotted against age and coloured by sex. $R$- hierarchical complexity, $C$- clustering coefficient, $r$- assortativity, $V$- degree variance, $L$- characteristic path length, $K$- mean rich-club coefficient.

| Index | $R$ | $C$ | $V$ | $r$ | $L$ | $K$ |
|---|---|---|---|---|---|---|
| Age | 0.6521 (0.55) | **0.0014 (5.78)** | **0.0013 (5.83)** | 0.1860 (1.65) | **0.0027 (5.19)** | **0.0011 (5.96)** |
| Sex | 0.0562 (3.77) | **0.0014 (11.02)** | **0.0002 (15.57)** | 0.1009 (2.76) | **0.0078 (7.49)** | 0.0527 (3.88) |
| Interaction | 0.7673 (0.38) | 0.3218 (1.18) | 0.5995 (0.63) | 0.0818 (2.33) | 0.4179 (0.96) | 0.1988 (1.59) |

We also implemented 2-way anovas within the tier-based analyses. Again, no associations were found between hierarchical complexity within any tier and the age and sex of participants, Table 2.

*Table 2. Results (p-values) from 2-way ANOVA tests across age and sex of within-tier complexity*

| | $R$ of Tier | 1(t) | 1b | 2(t) | 2b | 3(t) | 3b | 4(t) | 4b |
|---|---|---|---|---|---|---|---|---|---|
| **4-Tier** | **Age** | 0.5218 | - | 0.3909 | - | 0.1973 | - | 0.1480 | - |
| | **Sex** | 0.2886 | - | 0.1386 | - | 0.2935 | - | 0.6545 | - |
| | **Interaction** | 0.9293 | - | 0.2979 | - | 0.9584 | - | 0.9242 | - |
| **8-Tier** | **Age** | 0.3272 | 0.2978 | 0.5583 | 0.1127 | 0.4948 | 0.0600 | 0.0946 | 0.9276 |
| | **Sex** | 0.7349 | 0.5431 | 0.7685 | 0.5944 | 0.4516 | 0.1228 | 0.3521 | 0.9452 |
| | **Interaction** | 0.9346 | 0.6914 | 0.0667 | 0.4105 | 0.9258 | 0.5192 | 0.9695 | 0.5741 |

### III. Consistency of regions within tiers

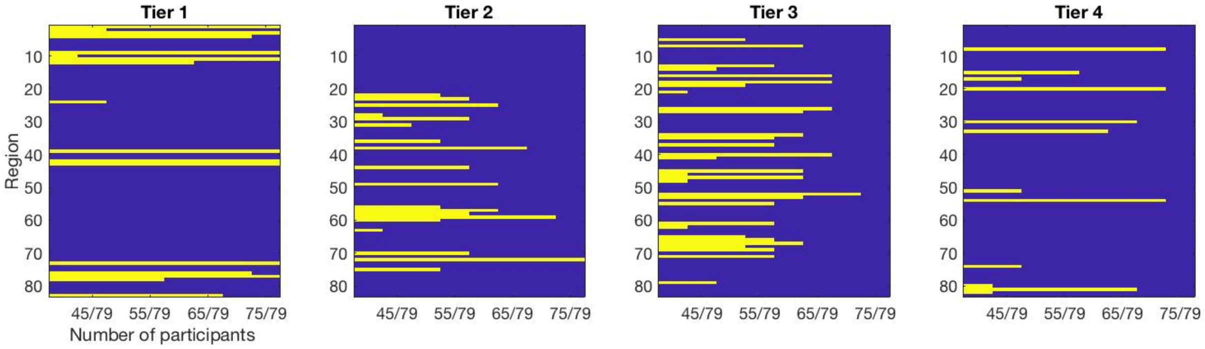


Figure 2. Consistency of regions within a given tier. Yellow indicates the region lies within the given tier for at least the number of participants denoted by the x-axis. Regions are enumerated as in Table 3. Axes labels as in the left-most plot.

In the main document we defined regions which were consistently within a single tier as one which appeared in that tier for at least two thirds of participants. Although this tells us those regions were in those tiers in the vast majority of cases, it is sensible to outline how the consistency of regions within tiers behaves if we were to consider stricter and more lenient thresholds. Therefore, tier categorisations were computed by changing the consistency level required, starting from 40/79 participants in steps of 5 up to 75/79 participants. The results are shown in Fig 6. In Fig 6 the enumerated regions are as per Table 3. Interestingly, the most consistent tiers were Tier 1 and Tier 4, which were found to be the least complex tiers in our document. Tiers 2 and 3 showed less regional consistency among participants.

Additionally, statistic of the mean degree within each Tier are as in Table 4.

*Table 3. Enumerations used for regions in the analysis. The order is of no significance.*

| | | |
|---|---|---|
| 1. Thalamus left | 29. Middle temporal gyrus left | 57. Inferior temporal gyrus right |
| 2. Caudate left | 30. Parahippocampal gyrus left | 58. Cingulate gyrus isthmus right |
| 3. Putamen left | 31. Paracentral gyrus left | 59. Lateral occipital cortex right |
| 4. Pallidum left | 32. Inferior frontal gyrus pars opercularis left | 60. Lateral orbitofrontal cortex right |
| 5. Brain stem | 33. Inferior frontal gyrus pars orbitalis left | 61. Lingual gyrus right |
| 6. Hippocampus left | 34. Inferior frontal gyrus pars triangularis left | 62. Medial orbitofrontal gyrus right |

| | | |
|---|---|---|
| 7. Amygdala left | 35. Pericalcarine cortex left | 63. Middle temporal gyrus right |
| 8. Accumbens left | 36. Postcentral gyrus left | 64. Parahippocampal gyrus right |
| 9. Thalamus right | 37. Posterior cingulate gyrus left | 65. Paracentral gyrus right |
| 10. Caudate right | 38. Precentral gyrus left | 66. Inferior frontal gyrus pars opercularis right |
| 11. Putamen right | 39. Precuneus left | 67. Inferior frontal gyrus pars orbitalis right' |
| 12. Pallidum right | 40. Rostral anterior cingulate cortex left | 68. Inferior frontal gyrus pars triangularis right |
| 13. Hippocampus right | 41. Rostral middle frontal gyrus left | 69. Pericalcarine cortex right |
| 14. Amygdala right | 42. Superior frontal gyrus left | 70. Postcentral gyrus right |
| 15. Accumbens right | 43. Superior parietal gyrus left | 71. Posterior cingulate gyrus right |
| 16. Banks superior temporal sulcus left | 44. Superior temporal gyrus left | 72. Precentral gyrus right |
| 17. Caudal anterior cingulate cortex left | 45. Supramarginal gyrus left | 73. Precuneus right |
| 18. Caudal middle frontal gyrus left | 46. Frontal pole left | 74. Rostral anterior cingulate cortex right |
| 19. Cuneus left | 47. Temporal pole left | 75. Rostral middle frontal gyrus right |
| 20. Entorhinal cortex left | 48. Transverse temporal gyrus left | 76. Superior frontal gyrus right |
| 21. Fusiform gyrus left | 49. Insula left | 77. Superior parietal gyrus right |
| 22. Inferior parietal gyrus left | 50. Banks superior temporal sulcus right | 78. Superior temporal gyrus right |
| 23. Inferior temporal gyrus left | 51. Caudal anterior cingulate cortex right | 79. Supramarginal gyrus right |
| 24. Cingulate gyrus isthmus left | 52. Caudal middle frontal gyrus right | 80. Frontal pole right |
| 25. Lateral occipital cortex left | 53. Cuneus right | 81. Temporal pole right |
| 26. Lateral orbitofrontal cortex left | 54. Entorhinal cortex right | 82. Transverse temporal gyrus right |
| 27. Lingual gyrus left | 55. Fusiform gyrus right | 83. Insula right |
| 28. Medial orbitofrontal gyrus left | 56. Inferior parietal gyrus right | |

*Table 4. Mean and standard deviation over participants of the average Tier degree*

| Tier | Tier 1 | | Tier 2 | | Tier 3 | | Tier 4 | |
|---|---|---|---|---|---|---|---|---|
| **Sex** | **Men** | **Women** | **Men** | **Women** | **Men** | **Women** | **Men** | **Women** |
| **Under 35** | 46.16 (3.14) | 50.72 (4.45) | 31.30 (2.44) | 33.37 (3.49) | 19.49 (1.88) | 21.37 (2.00) | 9.00 (0.86) | 10.25 (0.95) |
| **35 to 45** | 49.11 (4.35) | 51.18 (3.61) | 32.19 (2.92) | 34.24 (2.80) | 20.26 (2.65) | 20.59 (2.04) | 10.30 (1.71) | 11.05 (2.20) |
| **45 to 55** | 50.74 (5.66) | 51.87 (3.66) | 34.05 (4.22) | 35.23 (3.73) | 22.30 (3.61) | 21.69 (1.88) | 11.38 (3.04) | 10.46 (1.70) |
| **Over 55** | 52.10 (2.91) | 55.85 (2.47) | 35.05 (2.71) | 39.28 (2.90) | 22.08 (2.18) | 24.27 (2.66) | 10.31 (1.17) | 10.65 (0.83) |

### IV. Intensities of regional neighbourhood variance

In our article, we computed the neighbourhood degree variance of each region in each tier in order to understand more about the local regional patterns and how they may be reconciled with known neurophysiological. The mean values for each region over participants is shown in Fig 7 alongside values for configuration models. It is clear that values of configuration models appear to be independent of degree since all such values hover around the same value while the brain regions show variabilities which are consistent across participants and thus indicative of clear neurophysiological purpose for these values, whether high or low. The regions are enumerated as per Table 4.

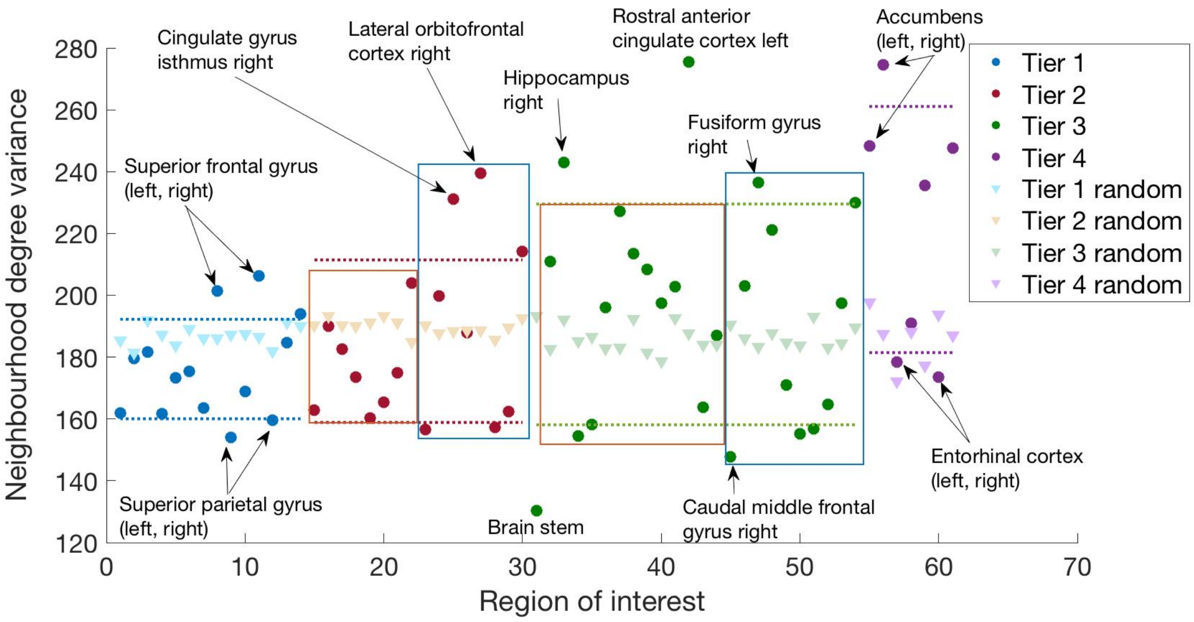


*Figure 7. Neighbourhood degree variance of Tier ROIs for structural connectomes and their random configuration models (random)*

*Table 5. Regions consistently within hierarchical tiers of the adult human structural connectome. Blue values are in Tier 1, orange in Tier 2, green in Tier 3 and purple in Tier 4.*

| | | |
|---|---|---|
| 1. Thalamus left | 22. Insula left | 43. Supramarginal gyrus left |
| 2. Putamen left | 23. Inferior parietal gyrus right | 44. Temporal pole left |
| 3. Pallidum left | 24. Inferior temporal gyrus right | 45. Caudal middle frontal gyrus right |
| 4. Thalamus right | 25. Cingulate gyrus isthmus right | 46. Cuneus right |
| 5. Putamen right | 26. Lateral occipital cortex right | 47. Fusiform gyrus right |
| 6. Pallidum right | 27. Lateral orbitofrontal cortex right | 48. Lingual gyrus right |
| 7. Precuneus left | 28. Postcentral gyrus right | 49. Paracentral gyrus right |
| 8. Superior frontal gyrus left | 29. Precentral gyrus right | 50. Inferior frontal gyrus pars opercularis right |
| 9. Superior parietal gyrus left | 30. Rostral middle frontal gyrus right | 51. Inferior frontal gyrus pars orbitalis right |
| 10. Precuneus right | 31. Brain stem | 52. Inferior frontal gyrus pars triangularis right |
| 11. Superior frontal gyrus right | 32. Amygdala left | 53. Pericalcarine cortex right |
| 12. Superior parietal gyrus right | 33. Hippocampus right | 54. Posterior cingulate gyrus right |
| 13. Superior temporal gyrus right | 34. Banks superior temporal sulcus left | 55. Accumbens left |
| 14. Insula right | 35. Caudal middle frontal gyrus left | 56. Accumbens right |
| 15. Inferior parietal gyrus left | 36. Cuneus left | 57. Entorhinal cortex left |
| 16. Inferior temporal gyrus left | 37. Lateral orbitofrontal cortex left | 58. Parahippocampal gyrus left |
| 17. Lateral occipital cortex left | 38. Lingual gyrus left | 59. Inferior frontal gyrus pars orbitalis left |
| 18. Middle temporal gyrus left | 39. Inferior frontal gyrus pars triangularis left | 60. Entorhinal cortex right |
| 19. Postcentral gyrus left | 40. Pericalcarine cortex left | 61. Temporal pole right |
| 20. Precentral gyrus left | 41. Posterior cingulate gyrus left | |
| 21. Superior temporal gyrus left | 42. Rostral anterior cingulate cortex left | |

Finally, the representations of the different tiers within notable ROI neighbourhoods were studied to elucidate specific connectivity patterns and see whether or not these could be explained by the known functional attributes of those ROIs. Effect sizes were computed using Cohen's $d$ between observed fractions of tier nodes and expected fractions of tier nodes of each subject, Table 6.

*Table 6. Effect sizes between the fractions of neighbouring nodes within each tier and their expected values for 15 ROIs*

| ROI | Tier 1 | Tier 2 | Tier 3 | Tier 4 |
|---|---|---|---|---|
| Superior Frontal Gyrus Left | 1.29 | 0.84 | -0.76 | -1.22 |
| Superior Frontal Gyrus Right | 1.46 | 0.68 | -0.85 | -1.30 |
| Superior Parietal Gyrus Left | 1.28 | 1.49 | -0.40$^{+}$ | -1.83 |
| Superior Parietal Gyrus Right | 1.15 | 1.44 | -0.32$^{+}$ | -1.80 |
| Cingulate Gyrus Isthmus Right | 1.69 | 0.36$^{+}$ | -1.44 | -1.38 |
| Lateral-Orbitofrontal Cortex Right | 1.26 | 0.16$^{+}$ | -1.09 | -0.32$^{+}$ |
| Brain stem | 1.74 | 1.29 | -1.78 | -1.90 |
| Hippocampus Right | 1.46 | 0.83 | -1.58 | -0.98 |
| Rostral Anterior Cingulate Left | 1.22 | -0.71 | -1.32 | 0.97 |
| Caudal Middle Frontal Gyrus Right | 1.69 | 1.28 | -1.56 | -1.89 |
| Fusiform Gyrus Right | 1.44 | 1.13 | -1.64 | -1.11 |
| Accumbens Left | 1.46 | -0.01$^{+}$ | -0.53 | -1.24 |
| Accumbens Right | 1.16 | 0.19$^{+}$ | -1.21 | 0.19$^{+}$ |
| Entorhinal Cortex Left | -0.32$^{+}$ | 0.22$^{+}$ | 0.62 | -0.94 |
| Entorhinal Cortex Right | -0.66 | -0.67 | 0.43 | 0.67 |

*Note: $^{+}$ indicates non-significant values at $\alpha$= 0.01 from a population t-test*
